# B cell transcriptomics reveals lasting dysregulation and rapid decline of protective immune memory after chronic hepatitis C cure

**DOI:** 10.1101/2025.10.24.664545

**Authors:** Fang Chen, Erick Giang, Padmaja Natarajan, Tony S. Mondala, Steven R. Head, Sing Chau Lau, Aishwarya Sundaresan, Liudmila Kulakova, Xiaohe Lin, Jasneet Aneja, Raiees Andrabi, Yun Zhang, Richard H. Scheuermann, Eric A. Toth, Thomas R. Fuerst, Lia Lewis-Ximenez, Arthur Y Kim, Paul Pockros, Georg M. Lauer, Mansun Law

## Abstract

Graphical abstract

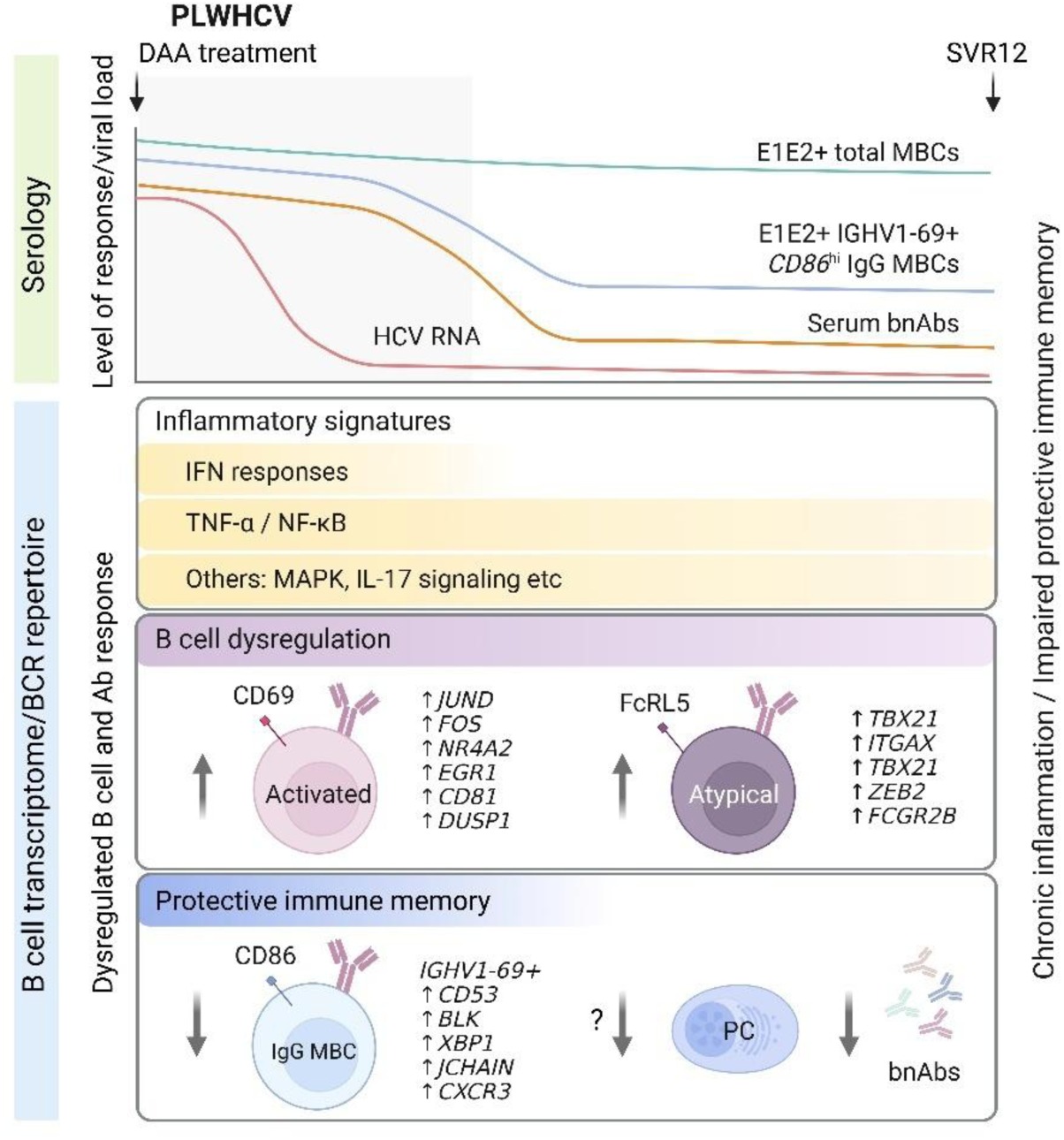

**Highlights:**

- B cells in CHC patients retained dysregulated transcriptional profiles despite successful DAA treatment
- B cell dysregulation is marked by global B cell hyperactivation and antigen-specific atypical MBC expansion
- Sustained upregulation of TNF-α signaling via NF-κB (TNF-α/NF-κB) is a central driver of persistent B cell dysregulation and chronic inflammation
- Viral clearance leads to restoration of IFN responses and IFN-stimulated gene (ISG) signatures in B cells but not TNF-α/NF-κB
- HCV IGHV1-69 bnAb-producing B cells are enriched within the *CD8*6^hi^ IgG MBC subset before treatment
- *CD86*^hi^ IgG MBC subset contracts rapidly post-cure in parallel with the rapid decline of cross-nAb responses and loss of protective immune memory against HCV reinfection

Chronic hepatitis C (CHC) disrupts host humoral immune response by impairing the timely generation of neutralizing antibody (nAb) and durable immune memory. However, the underlying mechanisms and their reversibility after viral clearance remain poorly defined. Here, through integrated single-cell transcriptomics and antibody repertoire characterization, we show that B cells from CHC patients retain transcriptional dysregulation even after successful antiviral therapy. Sustained TNF-α signaling emerged as a central driver of chronic B cell hyperactivation, persistent dysregulation and unresolved inflammation following cure. Furthermore, a *CD86*^hi^ memory B cell subset, responsible for an *IGHV1-69*-encoded multi-donor class recall nAb response, declined rapidly following viral clearance, compromising immune memory against reinfection. Together, these findings reveal how CHC imprints lasting B cell dysregulation, impairs nAb memory, and sustains inflammation in the B cell compartment, after viral clearance. The insights underscore the need for strategies aimed at restoring B cell homeostasis to achieve durable immune protection.

## INTRODUCTION

B cells play a central role in the humoral immune response against infections, primarily by producing antibodies, particularly neutralizing antibodies (nAbs), to eliminate pathogens. They also differentiate into long-lived plasma cells (LLPCs) and memory B cells (MBCs), providing durable immunity by sustaining antibody production and enabling rapid recall responses upon reinfection.^1^ In addition, some B cells can act as antigen-presenting cells (APCs) and regulatory B cells (Bregs), supporting cellular immune responses and the control of inflammation.^2^ Chronic infections, however, expose the host immune system to prolonged antigenic stimulation, leading to significant changes in the local immune microenvironment. These changes can directly or indirectly disrupt B cell-mediated immunity, resulting in altered B cell phenotypes and functions.^3,4^

Hepatitis C virus (HCV) infection progresses to chronic hepatitis C (CHC) in approximately 70% of those infected, greatly increasing their risk of life-threatening conditions such as cirrhosis and hepatocellular carcinoma (HCC).^5^ During CHC, the development of nAb responses is typically delayed,^6^ and virus-specific B cells undergo sustained activation and phenotypic reshaping, leading to expansion of B cell subsets such as activated MBCs (actMBCs) and atypical memory B cells (atyMBCs).^7–9^ AtyMBCs are characterized by expression of CD11c and T-bet, downregulation of CD27 and CD21, and diminished responsiveness to B cell receptor (BCR) stimulation.^10,11^ Expansion of these cells represents a hallmark of B cell dysregulation in chronic infections. Furthermore, the BCR repertoire in CHC patients is skewed, marked by biased gene usage and clonal expansion, predisposing individuals to autoimmune diseases and B cell-related malignancies.^7,12^

While direct-acting antivirals (DAAs) achieve sustained virologic response (SVR) in over 95% of CHC cases, they do not fully restore immune homeostasis.^13^ Viral clearance following successful DAA treatment leads to a rapid decline in serological antibody responses, particularly nAbs,^8,14^ which sharply contrasts with the long-lasting immunity typically observed for other viral infections or following live-virus vaccinations such as smallpox.^15^ This decline likely reflects a deficiency in LLPC formation or maintenance, raising concerns about the durability of protective immunity and vulnerability to HCV reinfection, particularly among high-risk populations where reinfection is frequently reported.^16,17^ Furthermore, DAA-induced viral clearance does not eliminate the risk of long-term complications.^18^ Some individuals recovered from CHC continue to exhibit signs of systemic and hepatic inflammation even after successful treatment.^19^ This persistent inflammation has been linked to an increased risk of tissue damage, including fibrosis and HCC, as well as extrahepatic complications such as lymphoproliferative disorders and cardiovascular disease.^19–21^ Dysregulated B cell responses may contribute to chronic inflammation through aberrant cytokine production, autoantibody secretion, and altered antigen presentation.^22^ In turn, the chronic inflammatory environment can impair humoral immunity by disrupting the generation and maintenance of LLPCs.^23–25^ Despite growing evidence of altered B cell and nAb responses in CHC before and after cure, the molecular mechanisms driving these changes remain poorly understood. Filling this knowledge gap will be important for devising strategies in preventing reinfection and improving vaccine responsiveness particularly in convalescent individuals. To address this, we set out to investigate B cell transcriptomics in people living with HCV (PLWHCV) undergoing DAA treatment.

We employed a recently developed soluble, secreted HCV E1E2 recombinant protein to isolate B cells responsible for producing nAbs.^26,27^ E1E2 constitutes the envelope glycoprotein complex on the viral surface and natural target of HCV nAbs, containing two major conserved neutralizing sites: (1) the antigenic region AR3 on the E2 neutralizing face (E2 NF),^28^ and (2) the E1-dependent bridging domain spanning E2 residues 646-704, which includes antigenic regions AR4 and AR5.^27,29,30^ Historically, studies of HCV-specific B cell and nAb responses relied on E2 subunit probes,^7–9,31^ due to the lack of natively folded E1E2 antigen. The recombinant E1E2 probe used here enables the isolation of B cells specific for both neutralizing sites. Notably, the majority of potent and broadly nAbs (bnAbs) against HCV identified to date target AR3 and are predominantly encoded by the human immunoglobulin heavy-chain variable gene *IGHV1-69*.^32,33^ These antibodies represent a multi-donor class of protective nAb responses detected not only in PLWHCV, but also in PLWHCV with potent and broad nAb responses known as elite neutralizers, and in recovered individuals resistant to multiple HCV reinfection.^33–35^

By integrating single-cell transcriptomics, BCR repertoire sequencing and antibody characterization, we determined the transcriptional and BCR repertoire changes of E1E2-specific B cells in CHC at pre-treatment (PreTx) and 12 weeks after the end of successful DAA treatment (SVR12). The results were compared to B cell responses to SARS-CoV-2 spike protein in vaccinated normal blood donors (NBDs). Our analysis revealed persistent transcriptional signatures of B cell hyperactivation and dysregulation in CHC even after viral clearance. Sustained upregulation of tumor necrosis factor alpha (TNF-α) signaling via nuclear factor-κB (NF-κB) appears to be a central driver of B cell dysregulation and contributes to chronic inflammation in the liver or other tissues, despite viral clearance. The chronic inflammatory state likely attenuates the durability of serological antibody memory,^23–25^ as evidenced by the rapid decline of circulating nAbs in CHC post-cure. Importantly, a *CD86*^hi^ MBC subset, responsible for an *IGHV1-69*-derived multi-donor class recall nAb responses, specifically contracted in parallel with viral clearance, leading to impaired protective immune memory in many recovered individuals. Together, these findings reveal how CHC imprints lasting B dysregulation, impairs nAb memory, and retains chronic inflammation in the B cell compartment post-cure, providing insights into strategies for restoring B cell homeostasis and durable immunity.

## RESULTS

### CHC-induced phenotypic and transcriptional alterations in B cell compartment persist after viral clearance

We first examined phenotypic alterations associated with CHC and DAA-mediated cure in E1E2-specific (E1E2^+^) and non-specific (E1E2^-^) B cells from the peripheral blood of 15 PLWHCV at PreTx and SVR12 (Table S1), using spectral flow cytometry (Figures S1 and S2). In comparison, spike-specific (spike^+^) and non-specific (spike^-^) B cells from 20 NBDs who had been recently vaccinated against SARS-CoV2 were also analyzed (Table S2), representing antigen-specific functional B cell responses and baseline healthy controls, respectively. On average, E1E2⁺ B cells comprised 0.17% (range: 0.06-0.4%) of total B cells in CHC, while spike⁺ B cells accounted for 0.34% (range: 0.1-0.8%) in NBDs (Figures S1B and S1C). The frequency of E1E2⁺ B cells remained stable in CHC before and after DAA cure (Figure 1C). Consistent with previous findings on E2⁺ B cells,^7–9^ E1E2⁺ B cells in CHC were marked by a persistent expansion of atyMBCs and actMBCs (Figures S1D-S1F), accompanied by elevated expression of multiple activation and dysregulation markers (Figure S1G). The activation marker CD69 was significantly increased across multiple subsets, particularly within naïve B cells (NBCs) (Figure S1H). Dysregulation-associated molecules, including FcRL5, T-bet, and CD11c, were preferentially expressed in atyMBCs, while PD-1 was elevated in both atyMBCs and actMBCs. Conversely, the liver-homing chemokine receptor CXCR3 was most abundant in actMBCs and CD21⁻ NBCs (Figure S1H). These phenotypic alterations persisted after viral clearance (Figures S1D-1G).

**Figure 1.**
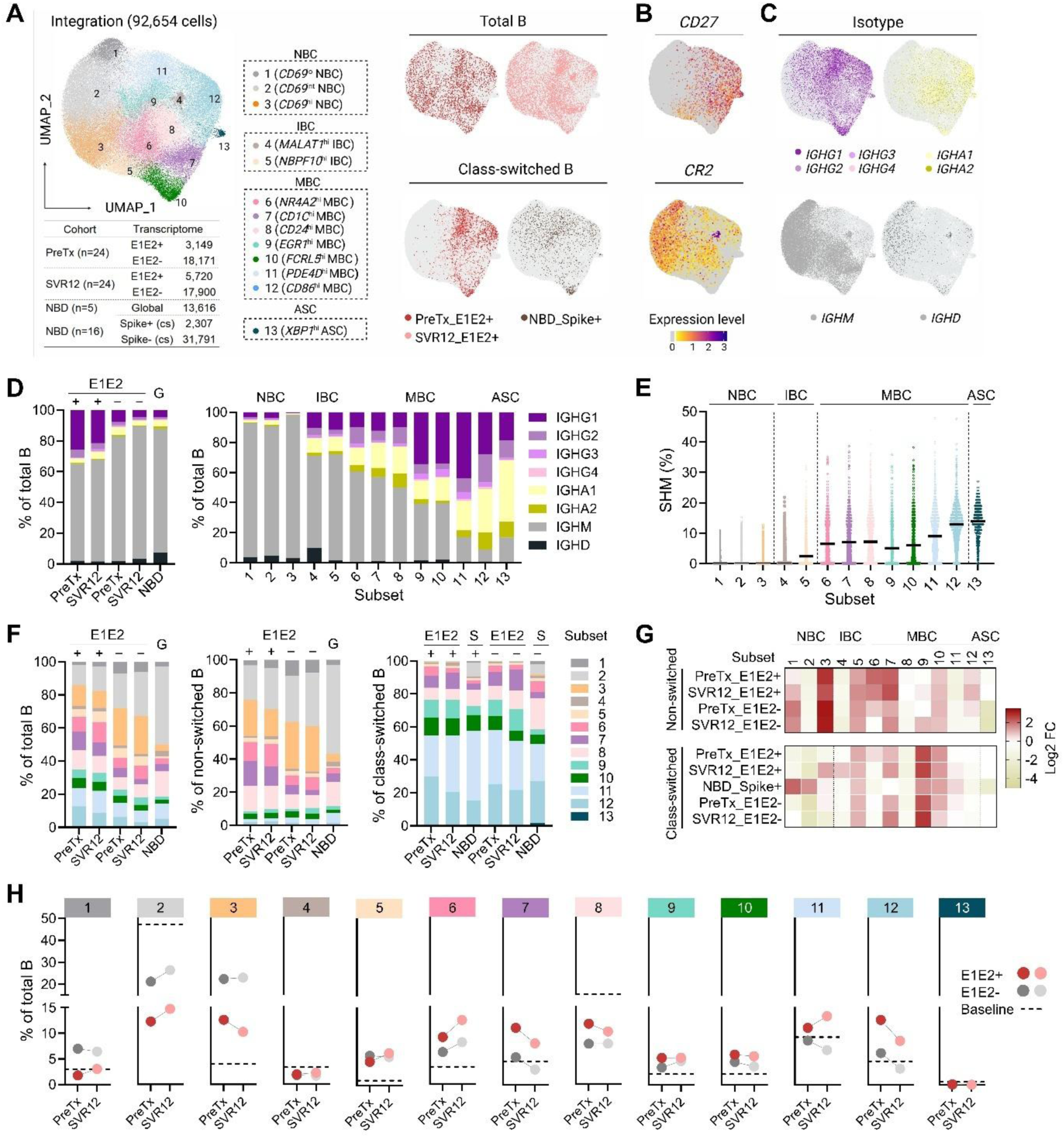
Single-cell transcriptomics reveals altered B cell composition in CHC before and after treatment. (A-C) Integrated uniform manifold approximation and projection (UMAP) of 92,645 B cells pooled from all samples. (A) UMAP visualization of B cell subsets across all samples (left), and antigen-specific B cells from CHC and NBDs (right). Subsets are grouped into 4 major populations: naïve B cells (NBC), intermediate B cells (IBC), memory B cells (MBC) and antibody-secreting cells (ASC). (B) Scaled gene expression of *CD27* and *CR2* (CD21). (C) UMAP projection of cells expressing different isotypes. cs, class-switched B cells. (D) Bar plots showing isotype distribution across cohorts and B cell subsets. G, global B cells. (E) Somatic hypermutation (SHM) frequency in B cells stratified by subset. Each dot represents a single cell. (F) Distribution of B cell subsets within total, non-switched and class-switched B cells across cohorts. G, global B cells; S, Spike. (G) Log2 fold change (FC) in B subset frequencies in CHC compared to baseline levels in NBDs. (H) Dynamic change in subset frequencies among E1E2^+^ and E1E2^-^ B cells in CHC before and after treatment. Dash lines represent the baseline levels in total B cells from NBDs.

To better define B cell heterogeneity and delineate how CHC and its resolution shape the transcriptional landscape, we conducted single-cell gene expression analysis of E1E2⁺ and E1E2⁻ B cells from 24 PLWHCV at PreTx and SVR12 (Table S1), together with global B cells from 5 NBDs and spike⁺/spike⁻ class-switched B cells from 16 NBDs (Table S2). After quality control and data integration, transcriptomes of 92,654 cells across all subjects and timepoints were obtained. Unbiased clustering of gene expression revealed 13 transcriptionally distinct B cell subsets (Figure 1A). Based on cell type-specific differentially expressed genes (DEGs; Figure S3) and previously documented B cell transcriptional profiles,^36–38^ these subsets were identified as NBCs (subsets 1-3), intermediate MBCs (IBCs, subsets 4 and 5), MBCs (subsets 6-12), and antibody-secreting cells (ASCs, subset 13). NBC subsets expressed high *CR2* (CD21) and low *CD27*, with predominantly IgM isotype and minimal somatic hypermutation (SHM) (Figures 1B-1E). While most MBC subsets upregulated both *CD27* and *CR2* (Figure 1B), subset 10 (*FCRL5*^hi^ MBC), showed downregulation of both markers, aligning with features of atyMBCs. In contrast, subset 7 (*CD1C*^hi^ MBC) and a fraction of cells within subset 12 (*CD86*^hi^ MBC) exhibited high *CD27* with low *CR2*, indicative of actMBCs. The IBC, MBC and ASC subsets comprised of both class-switched and non-switched B cells, with elevated rates of SHM (Figures 1C-1E).

Despite inter-individual variation (Figure S4), the overall B cell subset composition in CHC was markedly distinct from that of NBDs, featured by notable expansions of subsets 3 (*CD69*^hi^ NBC), 5 (*NBPF10*^hi^ IBC), 6 (*NR4A2*^hi^ MBC) and 7 (*CD1C*^hi^ MBC) among non-switched B cells, as well as subsets 9 (*EGR1*^hi^ MBC) and 10 (*FCRL5*^hi^ MBC) among class-switched B cells (Figures 1F and 1G). Subsets 3 and 9 exhibited the most pronounced increases (>5-fold compared to NBDs). Although these perturbations partially resolved following viral clearance at SVR12, they did not return to baseline (Figure 1H). Collectively, these findings highlight the durable impact of CHC on B cell composition, particularly within antigen-specific populations, even after viral clearance.

### Transcriptional signatures associated with B cell hyperactivation and dysregulation remain evident in CHC convalescents

To dissect the transcriptional mechanisms driving B cell dysregulation in CHC, we profiled gene signatures related to B cell activation, differentiation, and proliferation in both E1E2⁺ and E1E2⁻ B cells and mapped the contributing cellular subsets. At PreTx, most (60-80%) E1E2⁺ and E1E2⁻ B cells exhibited broad upregulation of activation-related genes relative to NBDs (Figures 2A left panel). These included *CD69*, *CD83*, members of the activator protein-1 (AP-1) family (e.g., *FOS*, *FOSB*, *JUN*, *JUNB* and *JUND*), members of the dual-specificity phosphatases (e.g., *DUSP1* and *DUSP2*), members of the nuclear receptor 4A family (e.g., *NR4A2* and *NR4A1*) and the tetraspanin *CD81*, with *JUND* being the most significantly upregulated (Figure 2E). The activation signatures were most prominent in subsets 3, 4, 5, 6, 9 and 10 (Figure 2A, right panel). The pronounced activation of NBCs (subset 3) aligns with previous reports^39^ and likely results from sustained stimulation through E2 interaction with CD81, a key HCV entry receptor expressed on hepatocytes as well as immune cells including B cells.^40^ After treatment, B cells from CHC retained elevated activation signatures compared to NBDs (Figure 2A, left panel). Key DEGs, such as *JUND*, *EGR1*, and *CD81*, remained significantly upregulated at SVR12 (Figure 2F).

**Figure 2.**
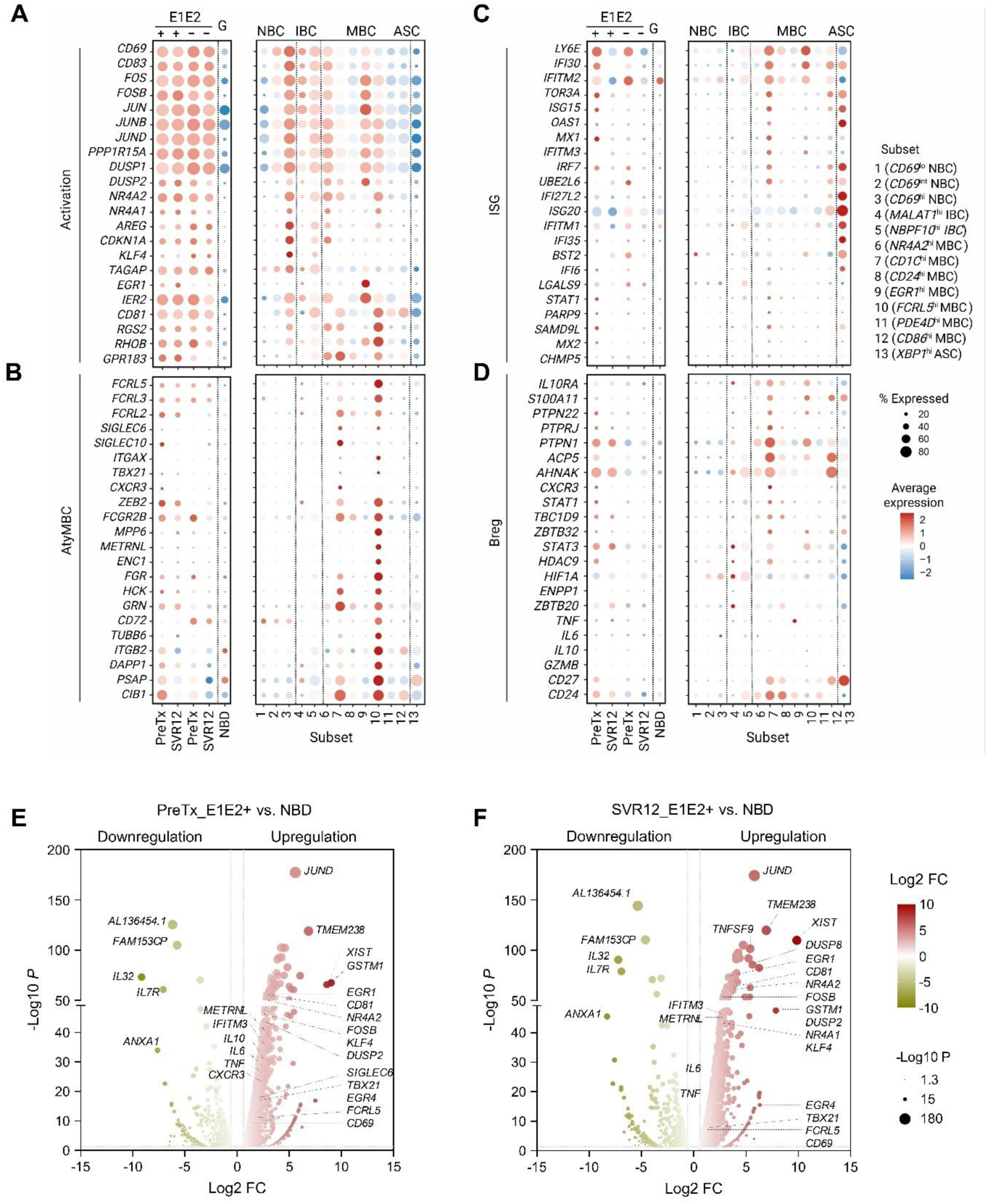
Gene signatures associated with B cell hyperactivation and dysregulation in CHC. (A-D) Dot plots showing expression levels of representative genes associated with B cell activation (A), atyMBCs (B), interferon-stimulated genes (ISGs) (C) and Bregs (D) in antigen-specific and non-specific B cells from CHC before and after treatment, compared to those in global B cells from NBDs. Corresponding gene expression levels in each B cell subset from CHC are also shown on the right. G, global B cells. (E-F) Volcano plots depicting DEGs in E1E2^+^ B cells from CHC, before (E) and after (F) treatment, compared to global B cells from NBDs. Dotted lines on the X-axis and Y-axis indicate the cutoff values for Log2 FC (|0.6|) and -Log10 *p*-value (1.3), respectively.

We also observed elevated expression of atyMBC-related genes^36,41,42^ in B cells from CHC patients (Figure 2B). In contrast to global B cell activation, these genes, such as *FCRL5, FCGR2B*, *SIGLEC10, TBX21* (T-bet), *ITGAX* (CD11c) and *ZEB2*, were predominantly upregulated in a small fraction (20-40%) of E1E2⁺ B cells (Figures 2B, left panel and 2E). Expression was highest in subset 10 (*FCRL5*^hi^ MBC) and, to a lesser extent, subset 7 (*CD1C*^hi^ MBC) (Figure 2B, right panel). After treatment, modest reductions were observed for a subset of these genes (e.g., *CXCR3*, *FGR* and *ITGB2*), while others (e.g., *FCRL5* and *TBX21*) remained significantly elevated (Figures 2B left panel and 2F), indicating partial but incomplete resolution of B cell dysregulation. Together, these findings provide transcriptional evidence that global B cell hyperactivation and E1E2⁺ B cell dysregulation in CHC persist despite viral clearance, indicating incomplete restoration of B cell homeostasis.

### CHC-induced ISG signatures in B cells are reversed after viral clearance

Given the importance of interferon (IFN) signaling in the regulation of inflammation in CHC,^43,44^ we analyzed IFN-related gene signatures across B cell subsets. Before treatment, a small population (∼20%) of E1E2⁺ and E1E2⁻ B cells exhibited increased expression of IFN-stimulated genes (ISGs) at PreTx (Figure 2C, left panel). Subset 7 (*CD1C*^hi^ MBC) was enriched for *LY6E*, *IFI30*, *IFITM2*, *ISG15* and *MX1*, whereas subset 13 (*XBP1*^hi^ ASC) displayed a distinct ISG profile marked by strong upregulation of *ISG20*, *IFI27L2*, and *IRF7* (Figure 2C, right panel). Notably, only a limited number of ASCs were recovered from CHC samples, likely due to their known sensitivity to cryopreservation.^45^ A restricted ISG signature was also observed in atyMBCs (subset 10), with elevated expression of *LY6E* and *IFI30* (Figure 2C, right panel). These results are consistent with previous reports of IFN signatures in specific B cell subsets.^46,47^ Following treatment, ISG expression (*IFI44L*, *IFI44*, *MX1*, *IFIT3*, *ISG15*) was markedly reduced at SVR12, restoring IFN-related transcriptional profiles to levels observed in NBDs (Figure 2C, left panel).

Intriguingly, subset 7 also displayed a CD24^hi^CD27^+^ phenotype (Figure 2D), indicative of a Breg subset.^48,49^ Supporting this, multiple genes associated with Breg differentiation,^50,51^ including *IL10RA*, *S100A11*, *ACP5* and *PTPN22*, were also upregulated in these cells (Figure 2D). Notably, increased expression of these genes was restricted to E1E2⁺ B cells in CHC and persisted after treatment (Figure 2D, left panel). In summary, IFN-driven transcriptional programs were upregulated in both E1E2⁺ and E1E2⁻ B cells during chronic infection and returned to baseline levels following viral clearance, in agreement with the successful elimination of viral infection.

### Persistent upregulation of TNF-α signaling sustains B cell dysregulation and chronic inflammation in CHC even after viral clearance

To better understand how altered transcriptional programs affect B cell functionality in CHC, we performed gene set and pathway analyses using the iPathwayGuide platform^52^ (Advaita Bioinformatics). Before treatment, we identified 3,677 DEGs (fold change >|1.5|) in E1E2⁺ B cells and 2,500 in E1E2⁻ B cells from CHC patients compared with global B cells from NBDs, of which 1,515 and 1,157 genes, respectively, exhibited fold change >|3| (Figures S5A, S6A and 3A). Most DEGs were upregulated, and enriched in pathways related to inflammation, cellular stress, and immune regulation (Figures S6A and S6B). Prominent hallmark signatures included TNF-α signaling via NF-κB (TNF-α/NF-κB), IL2-STAT5 signaling, TGF-β signaling, apoptosis, and interferon responses. Among these, TNF-α/NF-κB signaling emerged as the most significantly upregulated pathway (Figure S6B), consistent with its central role in chronic inflammation and immune activation.

After viral clearance, 3,713 and 2,728 DEGs were detected in E1E2⁺ and E1E2⁻ B cells from CHC compared to NBDs, respectively, with 1,467 and 922 genes showing fold change >|3| (Figures S5A, S6A, and 3A). The overall pathway profiles remained largely unchanged, suggesting that B cell transcriptional reprogramming is not fully reversed following viral clearance (Figures S6B, 3B and 3C). Notably, while TNF-α/NF-κB signaling, IL-2-STAT5 signaling, and MAPK pathways were highly enriched or affected in both E1E2⁺ and E1E2^-^ B cells, Wnt signaling and FoxO signaling pathways were primarily associated with E1E2⁺ B cells, whereas pathways related to viral protein-cytokine interaction and chemokine signaling were more pronounced in E1E2⁻ B cells (Figures 3B and 3C).

**Figure 3.**
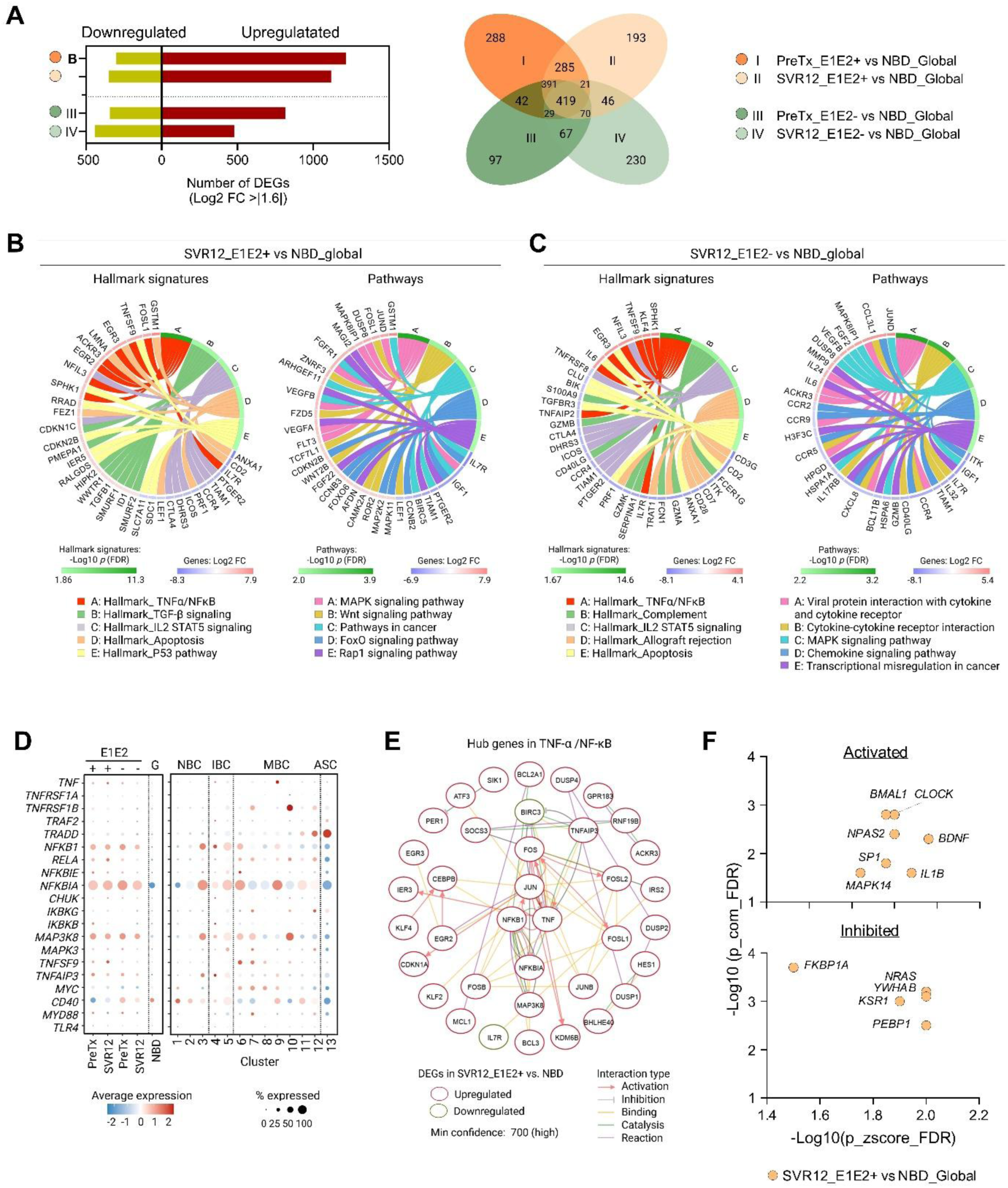
Altered pathways in B cells from CHC before and after treatment. (A) Numbers of DEGs with fold change (FC) > |3| (log₂ FC > |1.6|) in E1E2⁺ or E1E2⁻ B cells from CHC patients, before and after treatment, compared with global B cells from NBDs. A Venn diagram summarizing the meta-analysis was shown on the right. (B and C) Chord plots depicting significantly altered hallmark gene sets (MSigDB) and KEGG pathways, with representative DEGs, in E1E2⁺ (B) and E1E2⁻ (C) B cells from CHC patients at SVR12 relative to global B cells from NBDs. Statistical significances were assessed using false discovery rate (FDR)-adjusted *p*-values. (D) Expression of key genes associated with TNF-α signaling pathways in B cells and B cell subsets from CHC and NBDs. G, global B cells. (E) Network analysis of DEGs in SVR12_E1E2^+^ vs NBD_global showing the hub genes involved in TNF-α/NF-κB pathway. (F) Upstream regulators predicted to be activated (upper panel) or inhibited (lower panel) that target DEGs in E1E2^+^ B cells from CHC at SVR12 compared to global B cells from NBDs.

Furthermore, a direct comparison between CHC B cells before and after treatment revealed 95 DEGs in E1E2⁺ B cells and 127 DEGs in E1E2⁻ B cells at SVR12 (Figures S5B and S6A). These changes coincided with a significant reduction in IFN-α and IFN-γ responses (Figure S6B), as well as ISGs (e.g., *IFITM3*, *MX1* and *IFI44L*; Figure 2C), at SVR12. Importantly, these IFN-related signatures at SVR12 were no longer significantly different from those in NBDs (Figure S6B), suggesting that IFN signaling was fully restored following viral clearance. In contrast, although TNF-α/NF-κB and apoptosis-related signatures were partially downregulated in E1E2⁻ B cells at SVR12, their activity remained elevated relative to NBDs (Figure S6B), indicating incomplete resolution of inflammation and lingering transcriptional imprinting. Indeed, TNF-α/NF-κB remained the most significantly upregulated pathway at SVR12 (Figures 3B and 3C). Expression of key mediators and hub genes within this axis, including *FOS*, *JUN*, *NFKB1*, *TNF*, *NFKBIA*, and *MAP3K8*, remained highly upregulated in E1E2⁺ B cells even after viral clearance (Figures 2A, 3D and 3E).

In addition to persistent TNF-α/NF-κB, other inflammation-related signatures, such as complement, MAPK signaling, IL-17 signaling, C-type lectin receptor signaling, and AGE-RAGE signaling, were also significantly elevated in B cells from CHC, particularly within the E1E2⁺ subset (Figure S6C). Together, these sustained signaling perturbations reflect a persistent inflammatory program and a durable pro-inflammatory imprint in CHC convalescence.

To further investigate transcriptional regulators with the potential to restore B cell immune function in recovered individuals, we performed upstream regulator analysis on DEGs in E1E2⁺ B cells from CHC at SVR12 compared to NBDs. Several regulators predicted to be activated (e.g., *MAPK14* and *IL1B*) or inhibited (e.g., *FKBP1A* and *NRAS*) were identified (Figure 3F). Of these, *MAPK14* (encoding p38α, a MAPK family member) is of particular interest due to its interactions with transcription factors (e.g., *JUN*, *FOS*, *MEF2D* and *CEBPA*) and signaling molecules (e.g., *TNF* and *MAPKAPK2*) that regulate inflammation and stress responses (Figure S7), making it a promising target for therapeutic intervention in CHC post-cure.

### CHC infection and SARS-CoV2 vaccination induce atyMBCs with distinct transcriptional sub-profiles

Among MBCs, the atyMBC subset (subset 10, *FCRL5*^hi^ MBC) is notable for its unique transcriptional profile (Figure S3). It was distinguished from all other B cell subsets by 716 DEGs, with significant enrichment in hallmark signatures associated with TNF-α/NF-κB signaling, complement, and inflammatory response (Figures 4A and 4B). TNF-α/NF-κB was the most significantly upregulated pathway in this subset. Notably, *TNFRSF1B*, which encodes TNF receptor 2 (TNFR2), was specifically upregulated in atyMBCs (Figure 3D), implicating a TNFR2-mediated mechanism of TNF-α/NF-κB activation. This signaling axis is well-recognized for its roles in immune regulation, cell survival, and chronic inflammation,^53,54^ and may be a key driver of atyMBC dysfunction in CHC.

**Figure 4.**
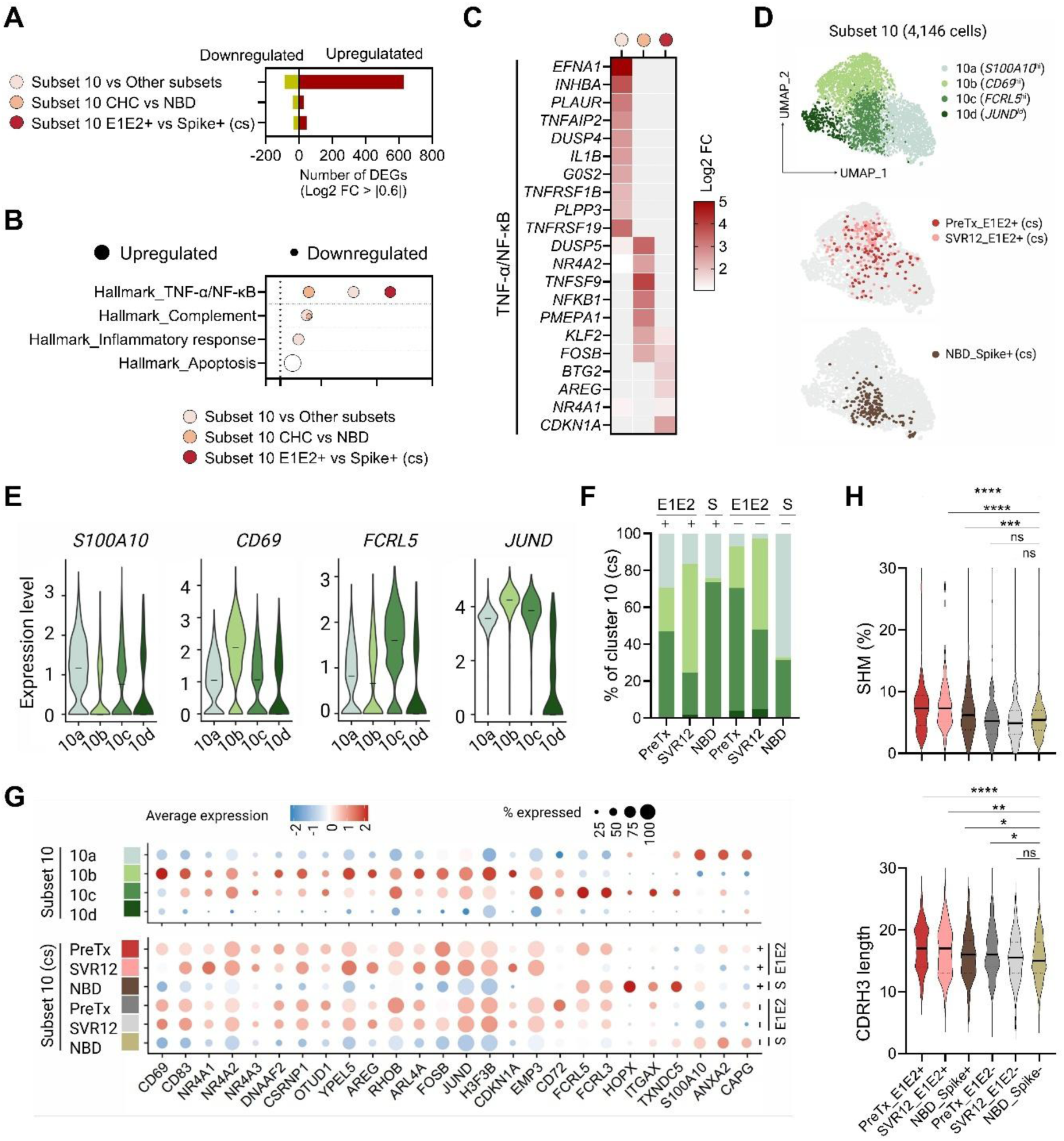
Transcriptional divergent atyMBCs (subset 10 cells) from CHC and NBDs. (A-C) DEG and Pathway analysis on the following comparisons: (1) Subset 10 vs other clusters, Cells from subset 10 versus from other subsets; (2) Subset 10 CHC vs NBD, subset 10 cells from CHC versus from NBDs; and (3) Subset 10 E1E2^+^ vs Spike^+^ (cs), subset 10 cells from class-switched E1E2^+^ B cells versus those from spike^+^ B cells. (A) Numbers of DEGs identified in each comparison. (B) Significantly altered hallmark signatures (MSigDB) in each comparison, with statistical significance assessed using FDR-adjusted *p*-values. Doted lines indicate the -Log10 (*p*-value) threshold of 1.3. (C) Log2 fold change (FC) of key DEGs associated with TNF-α/NF-κB for each comparison. (D) UMAP visualization of re-clustered subset 10 cells from CHC and NBDs. cs, class-switched B cells. (E) Expression levels of key genes differentially expressed in the subclusters of atyMBCs. (F) Distribution of atyMBC subclusters in class-switched (cs) B cells from CHC and NBDs. (G) Gene signatures of atyMBC across subclusters (upper panel) and subject cohorts (down panel). cs, class-switched. (H) Distribution of SHM rate and CDRH3 length in class-switched atyMBCs across cohorts. S, spike. Statistical significance was determined using two-tailed Mann-Whitney U-tests. *, *p*-value <0.05; **, *p*-value <0.01; ***, *p*-value <0.001; ****, *p*-value <0.0001; ns, not significant.

Comparison of atyMBCs from CHC and NBDs identified 65 DEGs. In addition, analysis of class-switched, antigen-specific atyMBCs revealed 78 DEGs when comparing E1E2⁺ B cells from CHC with spike⁺ B cells from vaccinated NBDs (Figure 4A). In both comparisons, TNF-α/NF-κB signaling again emerged as the top upregulated pathway (Figure 4B). While this signaling was consistently activated, the specific DEG profiles varied across conditions (Figure 4C), reflecting context-specific modulation.

To further delineate the heterogeneity within atyMBCs, we re-clustered subset 10 from all samples and identified four distinct subclusters (Figures 4D and 4E): 10a (*S100A10^hi^)*, 10b *(CD69^hi^)*, 10c (*FCRL5^hi^)* and 10d *(JUND^lo^)*. The subcluster 10b displayed a pronounced activation profile marked by elevated expressions of *CD69*, *CD83*, *NR4A2*, *FOSB* and *JUND* (Figure 4G). Subcluster 10c was also activated but with enhanced levels of inhibitory receptor genes (*FCRL3* and *FCRL5*) and *ITGAX*. Conversely, subclusters 10a and 10d showed minimal activation: 10a overexpressed *S100A10* and *ANXA2*, which together form a complex that inhibits p38 MAPK activation and apoptosis;^55^ 10d had lowered expression of all these markers.

Notably, class-switched atyMBCs displayed strikingly different subcluster distributions between CHC-derived and SARS-CoV-2 vaccine-induced B cell populations (Figures 4D and 4F). In NBDs, class-switched atyMBCs were predominantly composed of subclusters 10a and 10c. In contrast, CHC infection promoted expansion of subclusters 10b and 10c in both E1E2⁺ and E1E2⁻ B cells at PreTx, with subcluster 10c returning to baseline levels in E1E2⁺ B cells after DAA-mediated cure, whereas subcluster 10b remained elevated in both populations (Figure 4F). By comparison, SARS-CoV-2 vaccination selectively expanded subcluster 10c within spike⁺ B cells (Figure 4F). Furthermore, E1E2⁺ class-switched atyMBCs from CHC exhibited significantly higher SHM levels and longer CDRH3 loops (Figure 4H). Together, these findings highlight the heterogeneous transcriptional landscapes and functions of atyMBCs in distinct clinical contexts. CHC-induced atyMBCs displayed a more activated phenotype and lower *FCRL3/5* expression compared to those from vaccinated individuals (Figure 4G), consistent with the overall hyperactivated state driven by chronic inflammation. TNF-α/NF-κB signaling likely plays a central role in shaping the atyMBC transcriptome, contributing to their persistent dysregulation and the functional divergence observed between CHC and NBDs.

### CHC leads to a convergent clonal expansion in E1E2-specific BCR repertoire

Next, we conducted single-cell BCR repertoire sequencing and recovered 86,396 cells with matched V(D)J and transcriptome profiles across all samples (Table S3). The E1E2⁺ BCR repertoire in CHC was characterized by biased usage of *IGHV1-69* in IgG⁺ and *IGHV3-23* in IgM⁺ and IgA⁺ B cells (Figures 5A, S8A and S8B). At PreTx, *IGHV1-69*-derived E1E2^+^ IgG B cells were mainly enriched within the MBC subsets 9 and 12, followed by 4, 10 and 11, where IgG1 was the dominant isotype (Figures 5B and S8C). After viral clearance, these cells, particularly those within subset 12 (*CD86*^hi^ MBC), diminished quickly, suggesting a rapid loss of *IGHV1-69*-derived IgG MBCs following viral clearance. In contrast, *IGHV3-23*-derived E1E2^+^ IgM B cells were concentrated in NBC subsets 1, 2 and 3, and their frequencies remained stable before and after treatment, with the exception of subset 1, which showed an approximately 17% decrease at SVR12 (Figure 5B).

**Figure 5.**
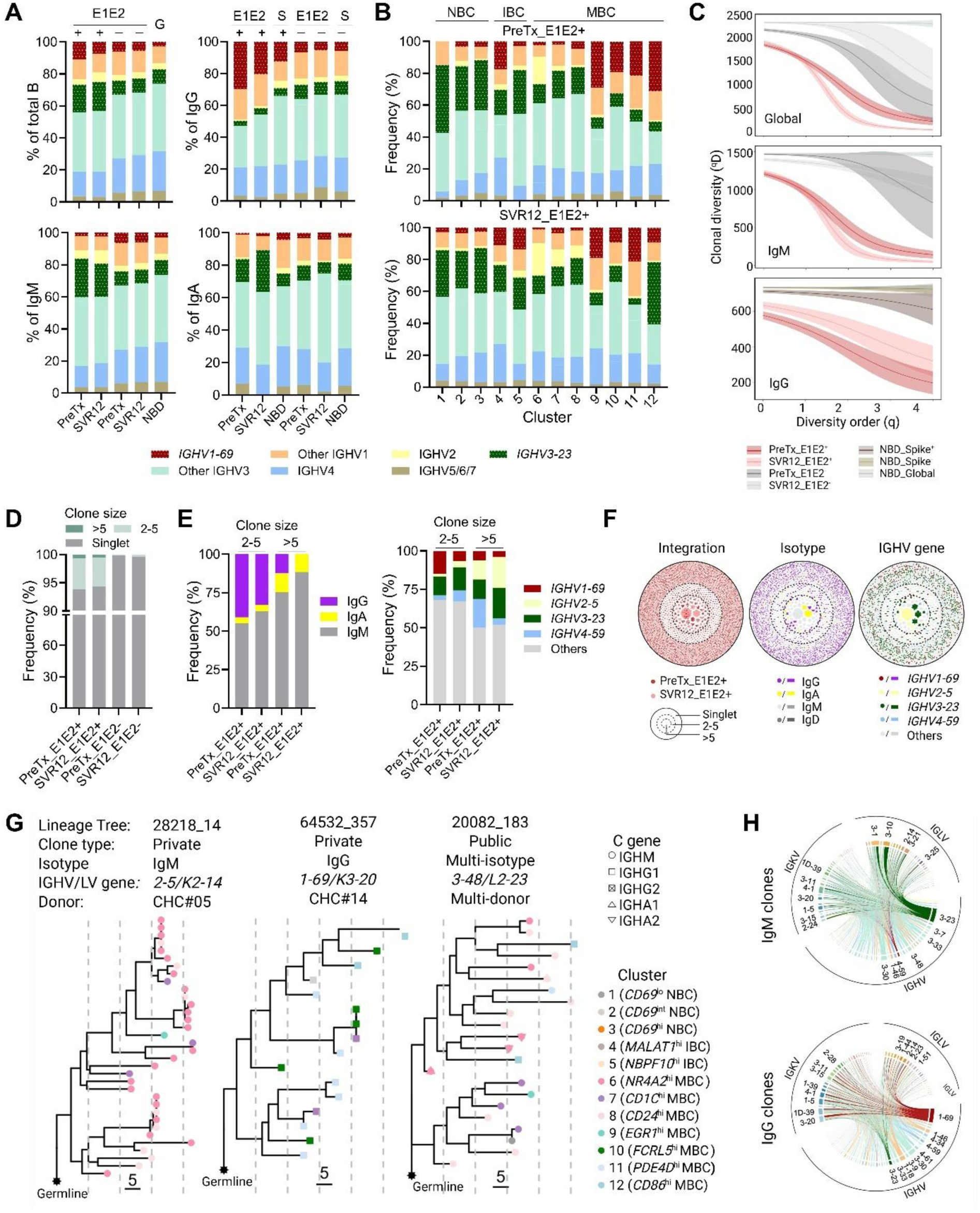
Antigen-specific BCR repertoires in CHC and NBDs. (A) Distribution of specific IGHV gene and gene family in the total, IgM, IgG and IgA repertoires across various cohorts. (B) Distribution of specific IGHV gene and gene family in E1E2^+^ B cells from CHC before and after treatment across B cell clusters. (C) Clonal diversity of BCR repertoires in CHC and NBDs, generated using Alakazam v.1.3.0. The diversity order (q) adjusts the weighting of clonotype frequencies, and clonal diversity (Hill number, qD) represents the effective number of distinct clonotypes in the repertoire. (D) Distribution of clonal families in E1E2^+^ and E1E2^-^ B cells from CHC before and after treatment. (E) Distribution of isotypes (left panel) and of the most frequently-used IGHV genes (right panel) in expanded clones (clone size>2) in E1E2^+^ B cells from CHC before and after treatment. (F) Clonal structure of integrated E1E2^+^ BCR repertoire of CHC before and after treatment. (G) Representative B cell lineage trees in the integrated E1E2^+^ BCR repertoire of CHC before and after treatment. (H) Circos plot showing paired heavy and light chain V genes in IgM- and IgG-expressing E1E2^+^ B cell clones.

The E1E2⁺ BCR repertoire in CHC also exhibited greater clonal expansion compared to their non-specific counterparts, along with reduced BCR diversity, indicative of a convergent antigen-driven B cell response (Figures 5C and S9A). The expanded clones in E1E2^+^ B cells were mostly small (clone size 2-5), and predominantly expressed unique isotypes, with IgM being the most prevalent, followed by IgG and IgA (Figures 5D and 5E). The most frequently utilized IGHV genes included *IGHV3-23*, *IGHV2-5*, *IGHV4-59*, and *IGHV1-69* (Figure 5E). Notably, *IGHV1-69*-derived IgG clones were predominant in small clonal families, while larger clonal families (clone size > 5) displayed a higher prevalence of *IGHV2-5*-derived IgA clones (Figure 5E).

Longitudinal tracking of E1E2^+^ B cell clones in CHC at PreTx and SVR12 showed that 68% (168 out of 269) of expanded clones persisted across time points, highlighting the presence of stable, enduring clones (Figure 5F). The majority of these clones were donor-specific (private clones), while a few were shared among multiple donors, representing a convergent public response at the population level (Figure S9B). Reconstruction of clonal lineage trees revealed that cells within a clonal family were derived from diverse B cell subsets (Figures 5G). Subsets 6 and 7 predominantly contained IgM clones, while subsets 10, 11, and 12 mainly consisted of IgG clones. Again, *IGHV3-23* and *IGHV1-69* were found to be enriched in the expanded IgM and IgG clones, respectively (Figure 5H). Overall, these findings point to a restricted and convergent BCR repertoire among E1E2-specific B cells, indicative of antigen-driven clonal selection.

### *IGHV1*-69-derived IgG MBC response in CHC declines rapidly after viral clearance

High-quality MBCs, especially those producing bnAbs, are key for preventing reinfection and maintaining protective immune memory. Given that the majority of AR3-targeting bnAbs originate from *IGHV1-69*,^32,33^ we investigated how CHC infection affects the E1E2^+^ IgG MBC response derived from this germline.

E1E2^+^ IGHV1-69 IgG MBCs in CHC exhibited specific genetic features, including (1) a long CDRH3 loop, averaging 19 amino acids compared to 16-17 in their non-specific counterparts; (2) elevated SHM rates (9% versus 6-8%); (3) favored Ig gene segments, such as *IGHD3-16*, *IGHD2-21*, *IGHJ6*, and *IGKV3-20*; and (4) increased usage of *IGHV1-69*01 and *10* gene alleles (Figures 6A-6D). The two *IGHV1-69* alleles correspond to the F and L allelic groups, respectively, which differ at a key polymorphic site in CDRH2 at position 54, where F alleles have phenylalanine (F) and L alleles have leucine (L) (Figure S10). Although both alleles have been found in bnAbs against HCV, influenza, HIV and SARS-Cov2,^32,56,57^ a recent immunization study using a humanized mouse model suggested that F alleles favor bnAb development, whereas L alleles are linked to polyreactive/autoreactive B cells that often become tolerized.^58^

**Figure 6.**
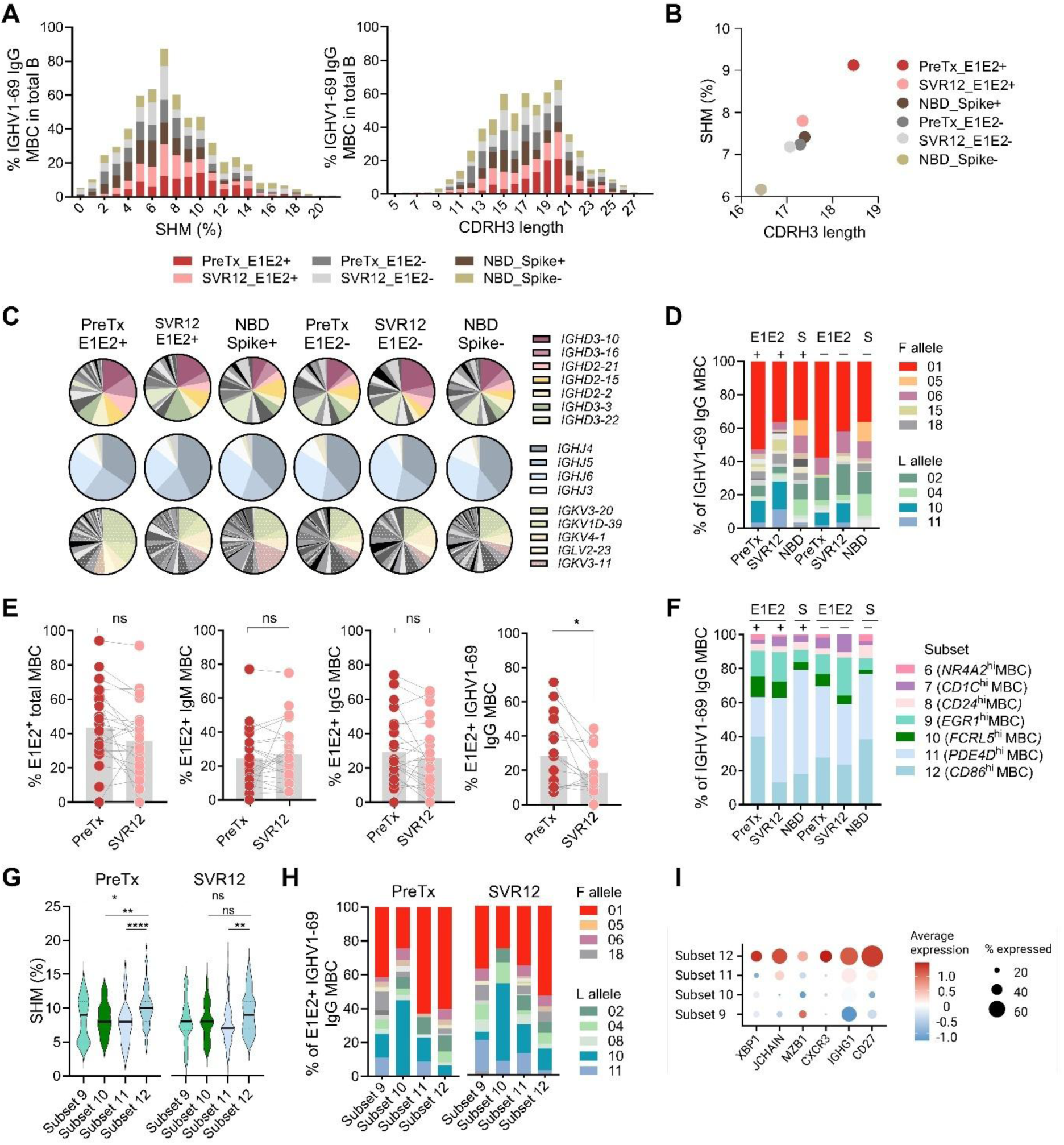
Molecular features of E1E2^+^ IGHV1-69 IgG MBCs in CHC. (A) Distribution of CDRH3 length and SHM rate in IGHV1-69 IgG MBCs from various cohorts. (B) Average CDRH3 length and SHM rate in IGHV1-69 IgG MBCs from various cohorts. (C) Distribution of heavy chain D and J gene segments, as well as light chain V gene segment in IGHV1-69 IgG MBCs from various cohorts. (D) Distribution of *IGHV1-69* gene alleles in IGHV1-69 IgG MBCs from various cohorts. (E) Frequency of total and subsets of E1E2^+^ MBCs in CHC before and after treatment. Each dot represents an individual patient. Statistical significance was determined using paired Wilcoxon t-tests. *, *p*-value <0.05; ns, not significant. (F) Distribution of IGHV1-69 IgG MBC subsets in various cohorts. Note the drop of cluster 12 MBC preTX versus SVR12. (G) SHM rate in E1E2^+^ IGHV1-69 IgG MBCs from subsets 9-12 in CHC before and after treatment. Statistical significance was determined using two-tailed Mann-Whitney U-tests. *, *p*-value <0.05; **, *p*-value <0.01; ****, *p*-value <0.0001; ns, not significant. (H) Distribution of *IGHV1-69* gene alleles in IGHV1-69 IgG MBCs from subsets 9-12 in CHC before and after treatment. (I) Expression of genes associated with antibody production in E1E2^+^ IGHV1-69 IgG MBCs from subsets 9-12.

E1E2⁺ IgG MBCs derived from *IGHV1-69* showed no major transcriptional differences compared to those utilizing other IGHV genes (data not shown). While the overall E1E2⁺ MBC frequencies remained steady post-treatment, IGHV1-69⁺ IgG MBCs significantly declined after viral clearance (Figure 6E), underpinning a specific loss of protective MBCs and bnAb memory against HCV soon after viral clearance. This decline was associated with diminished cross-neutralizing activity in plasma from CHC following viral clearance (Figure S11). Intriguingly, the contracted E1E2^+^ IGHV1-69 IgG MBCs were predominantly derived from subset 12 (*CD86*^hi^ MBC), which constituted the dominant IGHV1-69⁺ IgG MBC population at PreTx (Figure 6F). Following viral clearance, the frequency of subset 12 declined markedly, while subset 11 emerged as the predominant population.

E1E2^+^ IGHV1-69 IgG MBCs in subset 12 demonstrated elevated SHM, frequent usage of F alleles (especially *IGHV1-69*01*), and enhanced expression of antibody production-related genes such as *XBP1* (a crucial transcription factor essential for plasma cell differentiation^59^), *JCHAIN* (encodes the J chain, which is highly expressed in ASCs^60^) and *IGHG1*, as well as liver-homing chemokine receptor *CXCR3*, compared to those from subsets 9, 10 and 11 (Figures 6G-6I). These features suggest that the subset 12 MBCs are predisposed to differentiate into plasmablasts. In sum, our results highlight the potential role of E1E2^+^ IGHV1-69 IgG MBCs in bnAb production and memory response against HCV. These cells were predominantly present in the *CD86*^hi^ MBC subset (subset 12) and experienced a rapid decline after viral clearance, potentially weakening immune protection in individuals recovered from CHC.

### Functional analysis of antibody repertoires in CHC before and after viral clearance

To functionally evaluate E1E2⁺ B cells, we generated 157 monoclonal antibodies (mAbs) from paired Ig heavy and light chain sequences obtained from 22 individuals with CHC (Figure 7A). This panel included the most expanded clonotypes, antibodies with bnAb sequence features, and those lacking recognizable characteristics potentially novel HCV antibodies. All mAbs were expressed as IgG1 to enable direct functional comparison. Among them, 117 (75%) bound to H77 E1E2 in ELISA assays, including 52 derived from *IGHV1-69* and 65 using other IGHV genes (Figure 7A, left panel). Notably, 35 (67%) of the *IGHV1-69*-derived mAbs and 22 (34%) of those using other IGHV genes cross-neutralized three or more isolates from a panel of five HCV pseudoparticles (HCVpp) representing four tiers of neutralization resistance,^61^ and were thus classified as bnAbs (Figures 7A right panel and 7B).

**Figure 7.**
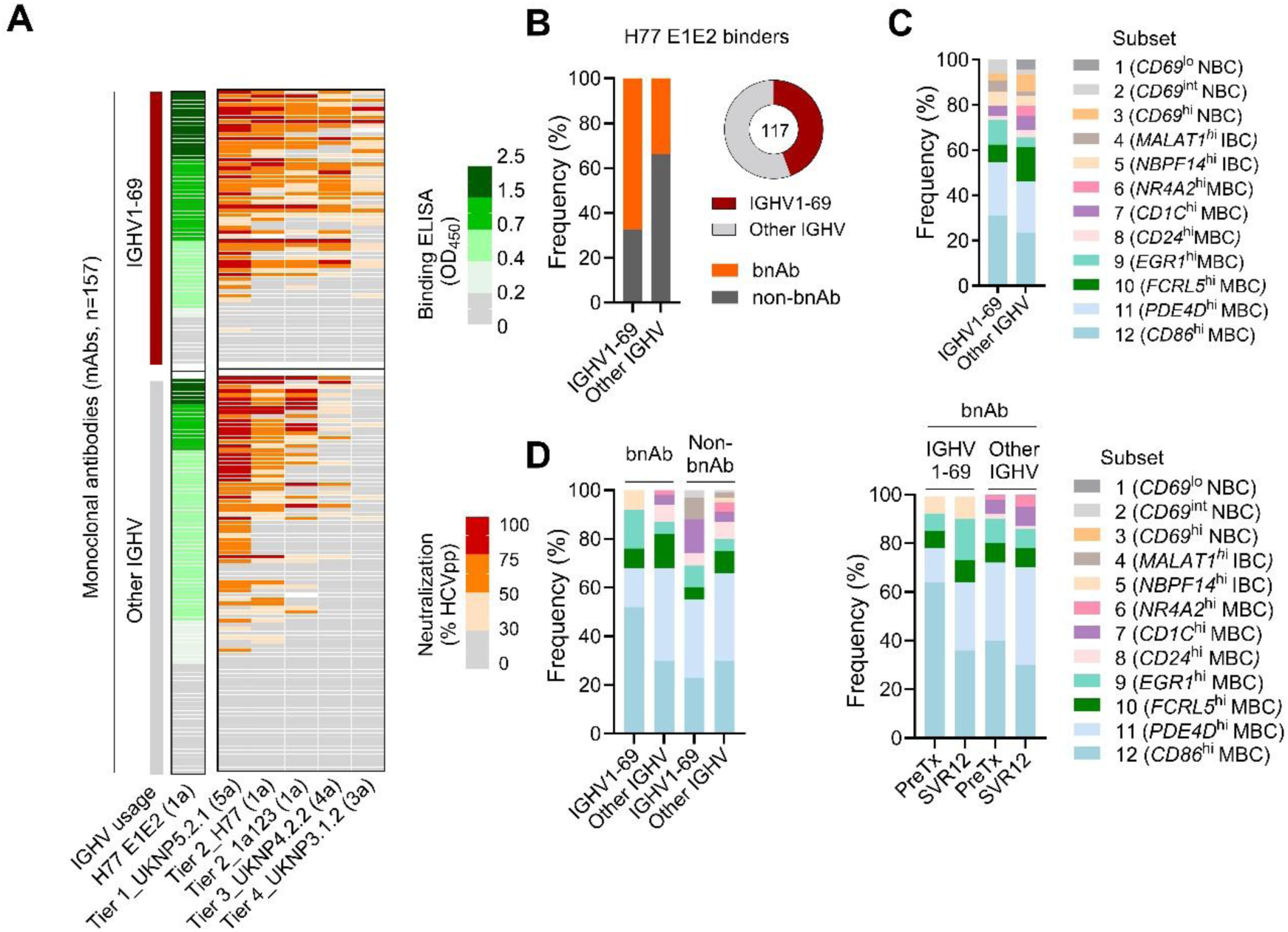
Functional properties and B cell subset origins of HCV bnAbs isolated from CHC. (A) Functional characterization of 157 mAbs generated from individuals with CHC by binding ELISA and HCVpp neutralization assays. (B) Distribution of bnAbs and non-bnAbs among *IGHV1-69*- and other IGHV-derived H77 E1E2-binding mAbs (n = 117). The relative proportions of IGHV1-69 and non-IGHV1-69 binders are shown to the right. (C) Distributions of B cell subsets among H77 E1E2-binding mAbs derived from *IGHV1-69* (n=52) versus other IGHV genes (n=65). (D) Left, distribution of B cell subsets among bnAbs and non-bnAbs derived from IGHV1-69 or other IGHV genes. Right, comparison of bnAb subset distributions in mAbs isolated before (PreTx) and after DAA-mediated viral clearance (SVR12).

Analysis of the origins of *IGHV1-69*- and other IGHV-derived HCV mAbs revealed overall similar B cell subset distributions, with subsets 12 (*CD86*^hi^ MBC) and 11 (*PDE4D*^hi^ MBC) representing the most prevalent populations (Figure 1C). However, when focusing on bnAbs, the subset composition differed significantly between the two groups: *IGHV1-69*-derived bnAbs were preferentially associated with subset 12, whereas bnAbs using other IGHV genes were mainly derived from subset 11. This differential pattern was not evident among non-bnAbs (Figure 7D, left panel). Longitudinal analysis showed a marked reduction of subset 12 in *IGHV1-69*-derived bnAbs after viral clearance, while the frequency subset 12 within other IGHV-derived bnAbs remained relatively stable (Figure 7D, right panel). Together, these findings confirm that E1E2⁺ B cells in CHC can generate potent cross-neutralizing antibodies, with *IGHV1-69*-derived *CD86*^hi^ IgG MBCs constituting a major source of this response. Importantly, HCV clearance is accompanied by the rapid and preferential loss of these bnAb-producing memory B cells, leading to loss of protective humoral memory.

## Discussion

HCV is currently the only chronic viral infection that can be completely eradicated with highly specific antiviral drugs, offering a unique model to study human immunology in the context of persistent infection, chronic inflammation, and immune restoration. In this study, we leveraged this model to examine how chronic infection and its resolution shape the antigen-specific B cell landscape and antibody repertoires, which are critical for recall responses to reinfection and vaccination. To achieve this, we integrated single-cell gene expression, BCR repertoire sequencing and antibody functional characterization, to examine the different B cell subsets during antiviral therapy. Here, we focus on the transcriptional states of B cell subsets following viral clearance. Detailed characterization of new bnAbs targeting conserved HCV neutralization sites is described in two accompanying manuscripts (in preparation).

We demonstrated that, while most individuals cleared HCV RNA by week 4 after DAA initiation,^62^ B cell dysregulation was incompletely reversed and persisted at SVR12. This ongoing B cell dysregulation is marked by sustained global B cell hyperactivation and persistent antigen-specific atyMBC expansion (Figures 1-4).

The sustained B cell hyperactivation and inflammation is particularly concerning, as it may result from unresolved hepatic or extrahepatic inflammation despite successful antiviral treatment, which may in turn contribute to the inflammatory state.^19,63^ Persistent low-grade hepatic inflammation, particularly in the portal regions, can occur in both native or transplanted liver following treatment and contribute to the development of cirrhosis and HCC in cured individuals.^64,65^ Our transcriptomic analysis revealed that multiple inflammation-related hallmark signatures and pathways, including TNF-α/NF-κB, complement, MAPK signaling and IL-17 signaling, remained persistently upregulated even after viral clearance, contributing to ongoing B cell hyperactivation in CHC convalescents (Figure S6). Among these, TNF-α/NF-κB signaling emerged as a central driver.

TNF-α is a major pro-inflammatory cytokine. It engages with two receptors, TNFR1 (*TNFRSF1A*) and TNFR2 (*TNFRSF1B*), triggering multiple downstream pathways, primarily the NF-κB and MAPK signaling cascades.^66^ These pathways coordinate key cellular processes including inflammation, cell survival, proliferation and differentiation, as well as apoptosis.^67^ They also promote the production of inflammatory mediators, including IL-1, IL-6, IL-8, and TNF-α itself, which can further amplify immune activation. Notably, TNF-α production is not strictly dependent on viral infection and may persist in the context of ongoing tissue damage, immune dysregulation, and fibrotic remodeling established over decades of chronic inflammation.^66,68^ Such fibrotic microenvironments may act as reservoirs for TNF-α and other pro-inflammatory cytokines, thereby promoting sustained B cell hyperactivation. The sustained hyperactivation and upregulated TNF-α signaling can suppress plasma cell production.^23–25^ This is evident in the rapid decline of circulating nAbs and the relevant MBCs following viral clearance (Figure S11).

Intriguingly, atyMBCs (subset 10) exhibited significantly elevated TNF-α/NF-κB signaling compared to other B cell subsets (Figure 4B), along with selective upregulation of *TNFRSF1B* (Figure 3D). These findings are in line with established roles of TNF receptors: TNFR1 is broadly expressed and linked to proinflammatory and apoptotic responses, whereas TNFR2 is more immune cell-specific and facilitates survival and immune regulation.^53,54^ Although the expansion of atyMBCs has been documented in a variety of conditions, such as chronic infections, autoimmune diseases, acute viral infections and vaccinations,^10,69–71^ their precise role across different immune stimuli and disease contexts remains unclear. Here, by comparing atyMBCs from individuals with CHC and SARS-CoV-2 vaccinated NBDs, we demonstrated that atyMBCs from CHC exhibit a heightened activation state, with activation signals primarily enriched in subset 10b (Figures 4F and 4G). This subset appears to be a hallmark of chronic infections including CHC, malaria, and HIV, but is absent in vaccine-induced responses.^41,42,72^ The hyperactivated state of atyMBCs may impair their ability to produce antibodies or differentiate into LLPCs, despite their retained potential to generate bnAbs against HCV (Figure 7D). CHC-induced atyMBCs remained expanded after viral clearance (Figure 1H), with only partial resolution of their dysregulated gene signatures (Figure 2B), suggesting that ongoing TNF-α/NF-κB signaling via TNFR2 plays a key role in maintaining their dysfunctional state (Figure 4B).

IFN responses also contribute to B cell dysregulation and inflammation. CHC induces a systemic type I IFN response, resulting in widespread expression of ISGs across immune cell populations.^73,74^ These type I IFNs (particularly IFN-α and IFN-β) help establish a global antiviral state, however, their sustained expression in chronic infection can dampen adaptive immunity, impair germinal center responses, and promote the expansion of ISG-expressing atypical B cells.^75,76^ Unlike TNF-α, type I IFNs are primarily induced by viral RNA recognition through innate immune sensors such as RIG-I, MDA5, and TLR7/9.^75^ Following HCV clearance, the stimulus for IFN production is eliminated, leading to a rapid decline in ISG expression in both peripheral blood and the liver, often within weeks after viral clearance. Supporting this, we found that ISG-expressing B cells were primarily enriched in subset 7 (*CD1C*^hi^ MBCs) of E1E2⁺ B cells, and these IFN-driven features were fully resolved at SVR12 (Figures 2C and S6B).

Dissecting E1E2-specific BCR repertoires from CHC revealed isotype-specific biases in IGHV germline usage and clonal expansion: *IGHV3-23* was preferentially utilized in IgM responses, while *IGHV1-69* dominated the IgG compartment (Figure 5). The majority of *IGHV3-23*-derived antibodies, with a few exceptions, recognize isolate-specific sites that HCV can readily escape.^7,30,77^ In contrast, most *IGHV1-69*-derived antibodies exhibited broad neutralizing activity (Figure 2B), representing an important multi-donor bnAb class that is conserved in both humans and rhesus macaques.^28,30,33–35,78–80^ The observed clonal expansion biases between these two MBC subpopulations suggest different selective pressure in the IgM and IgG isotype compartments for recognition of conserved viral sites of vulnerability. Tracking MBCs encoding for bnAbs are crucial for understanding cross-neutralizing immune memory against reinfection and vaccination. Furthermore, in this study, bnAbs of similar or even greater frequency and potency were identified in typical PLWHCV compared with HCV elite neutralizers^33^ or reinfection-resistant recovered individuals^78^ (Figure 7A), highlighting the importance of comprehensive BCR repertoire analysis to uncover additional potent and cross-reactive bnAbs that can guide vaccine design.

Following viral clearance, we observed an unusual rapid loss of IGHV1-69 IgG MBCs specifically in the *CD86*^hi^MBCs (subset 12) (Figures 5B and 6F). These cells are pivotal producers of HCV bnAbs (Figure 7). Their rapid decline is likely driven by the loss of ongoing antigen stimulation and inflammatory cues that previously sustained their activation and expansion, and may account for the short-lived circulating nAb responses observed after recovery (Figure S11). Whether these cells are transitioning to a more quiescent state or undergoing apoptosis due to metabolic stress, replicative exhaustion, or lack of survival signals remains to be determined. Nevertheless, the contraction of this critical MBC population could result in the loss of protective nAb memory, potentially compromising immunity against HCV reinfection. A recent study profiling the transcriptional state of E2-specific B cells reported an elevated inflammatory signals and suggested a potential link between MBC quality and protection against reinfection.^81^ Further investigation is needed to elucidate how persistent inflammatory programming in B cells affects humoral immunity, particularly the IGHV1-69 bnAb response, to HCV.

Taking together, our findings highlight that prolonged HCV exposure leads to lasting molecular alterations in the B cell compartment even after viral clearance. These alterations reflect and reinforce a persistent inflammatory milieu that hampers plasma cell differentiation and skews MBC composition, weakening protective immunity upon reinfection. The sustained inflammation may also contribute to ongoing tissue damage, exacerbating liver pathology and systemic complications despite viral clearance.^18,19^ Addressing this inflammation is critical for improving long-term health outcomes after HCV cure. Our transcriptomic analysis identifies *MAPK14* as a key immune regulator (Figure S7) with therapeutic potential to modulate inflammation. Inhibitors targeting *MAPK14* such as SB203580 have shown promise in reducing inflammation and tissue pathology in various disease models.^82–84^ Long-term follow-up and further investigation into the dysregulated signaling pathways and transcriptional regulators will be crucial for restoring B cell homeostasis and enhancing durable immunity in cured CHC and other immune-mediated diseases. Last but not least, with advances in single-cell technology and data analysis pipelines, the study of immune cells in easily accessible clinical samples, such as peripheral blood lymphocytes, enables highly informative assessments of immune status. B cells, in particular, have the potential to serve as sensitive “biosensors” for detecting inflammation and immune dysregulation. In the context of viral hepatitis, a controlled human infection model (CHIM) is actively being developed.^85–87^ As such, it is essential to monitor the effects of controlled infection on the immune system of volunteers using accessible clinical specimens and high-throughput approaches. Transcriptomic profiling^88^ will be instrumental in tracking both innate and adaptive immune responses and in determining whether systemic T cell exhaustion and B cell hyperactivation occur in response to the different infection conditions to be examined in CHIM.

## Methods

### Human samples

Longitudinal blood samples from 24 individuals with CHC undergoing direct-acting antiviral (DAA) treatment were collected at Massachusetts General Hospital (MGH) between 2012 and 2016 (clinical trial NCT02476617). Patient demographics and clinical characteristics are described in Table S1. Additionally, single-time point blood samples from 24 NBDs were collected at Scripps Research during the SARS-CoV-2 pandemic from March to May 2022. Peripheral blood mononuclear cells (PBMCs) and plasma were processed using Lymphoprep and SepMate tubes (StemCell) according to manufacturer’s instructions. Plasma samples were heat-inactivated at 56°C for 30 minutes, aliquoted and stored at −80°C. PBMCs were aliquoted and stored in liquid nitrogen. The vaccination and infection status for SARS-CoV-2 in NBDs was confirmed by detecting plasma antibodies against SARS-CoV-2 spike and nucleoprotein proteins through ELISA, and the results are detailed in Table S2.

### Antigen probes

To prepare biotinylated HCV sE1E2, a soluble, secreted E1E2 construct bearing a C-terminal Avi tag (sE1E2-Avi) from an H77 viral stain was transiently co-expressed with human furin in Expi293 cells using the Expi293 Expression System (ThermoFisher Scientific), according to the manufacturer’s protocols. Culture supernatants were harvested 72 hours post-transfection, clarified by centrifugation, and filtered through a 0.45 μM filter. The sE1E2-Avi protein was purified using sequential immobilized metal affinity chromatography and size exclusion chromatography, as previously described.^26,27,89^ Biotinylation was carried out using the BirA biotin-protein ligase kit (Avidity, Aurora, CO), following the manufacturer’s recommendations. The structural integrity of biotinylated sE1E2-Avi was confirmed by dose-response ELISA using a panel of conformation-specific monoclonal antibodies.^26^ Biotinylated SARS-CoV-2 spike protein was produced as described previously.^90^

### Spectral flow cytometry

To stain antigen-specific B cells, the biotinylated antigen probes were individually conjugated with a PE streptavidin (ThermoFisher Scientific) or a BV421 streptavidin (BioLegend) at a 4:1 molar ratio, as previously described.^91^ The probes were mixed in 1:1 Brilliant Buffer (BD Bioscience) and FACS buffer (PBS with 2% FBS and 2 mM EDTA) with 5 μM free D-biotin.

Cryopreserved PBMCs were thawed and first stained with LIVE/DEAD Fixable Aqua dead cell stain dye (ThermoFisher Scientific) in 96-well U-bottom plates to exclude dead cells. The cells were then blocked with an anti-CD81 antibody (5 μg/mL, BD Biosciences) and Fc blocker (TruStain FcX™, BioLegend) before being stained with biotinylated probes (200 ng E1E2 and 500 ng spike per color per 50 μL, respectively) at 4°C for 30 min. This was followed by an additional 30 minutes of staining with a cocktail of surface markers at 4°C in the dark. Samples were analyzed on a Cytek Aurora spectral flow cytometer equipped with five lasers, using the SpectroFlo software (Cytek Biosciences). Data analysis was conducted using FlowJo software (BD, version 10.9.0). The gating strategies employed are detailed in Figure S3.

### Single-cell RNA library preparation and sequencing

Antigen-specific and non-specific B cells were sorted on a MoFlo® Astrios EQ (Beckman Coulter) into 0.04% BSA in PBS. Sorted B cells were washed twice with PBS + 0.04% BSA and adjusted to a concentration of 1,000 cells/µL. They were then processed using the 10X Genomics workflow for gene expression (GEX) and BCR immune profiling [V(D)J], utilizing the Chromium Next GEM Single Cell 5’ HT Kit v2, Single Cell Human BCR Amplification Kit, and 5’ Feature Barcode Kit, according to the manufacturer’s instructions (10X Genomics).

Single-cell 5’ GEX and V(D)J sequencing data from pooled samples were generated on a NovaSeq6000 sequencer (Illumina) using 26 x 90 paired end reads with 10 base dual indexing. Raw FASTQ files were aligned to the GRCh38 reference genome [refdata-gex-GRCh38-2020-A and refdata-cellranger-vdj-GRCh38-alts-ensembl-5.0.0 for GEX and V(D)J, respectively].^92^ Count matrices for cell barcodes and UMIs were generated by Cell Ranger (version 6.0.0, 10X Genomics). The multi pipeline with default parameter settings was used for demultiplexing and quantifying gene expression and assembly of BCR sequences.

### Transcriptome analysis

Cell Ranger (version 6.0) output was analyzed in R (version 4.3.2) using Seurat 5.0 package. Cells expressing <200 or >3,000 genes, >10% mitochondrial genes, and <10% ribosomal protein genes were removed. To generate clusters that were uniformly present in all samples, the reciprocal principal component analysis (PCA) method was used for integrated clustering in Seurat (version 5). Data were normalized, transformed and variable genes were detected using default parameter settings. Uniform manifold approximation and projection (UMAP) was performed for low-dimensional embedding of the data, using SCTransform, RunPCA and RunUMAP with 30 principal components. Clusters with transcriptionally similar profiles were identified by shared nearest neighbor (SNN) and Louvain algorithm for modularity optimization, using FindNeighbors and FindClusters in PCA space with a resolution of 0.5.

Cell type-specific differentially expressed genes (DEGs), determined with default cutoffs on log2 fold change of 0.6 and an adjusted *p*-value < 0.05, were comprehensively analyzed for enriched gene sets (The Molecular Signatures Database, MSigDB), pathways (Kyoto Encyclopedia of Genes and Genomes, KEGG), and upstream regulators using the Advaita iPathwayGuide platform (https://www.advaitabio.com/ipathwayguide).

### BCR repertoire analysis

The single-cell BCR sequencing data were processed with 10X Genomics Cell Ranger vdj. 10X Genomics Loupe V(D)J Browser was used to visualize V(D)J sequences and clonotypes. Ig V(D)J annotation and C gene assignment was performed using IgBLAST v 1.22.0 (https://www.ncbi.nlm.nih.gov/igblast/). Change-O v.1.3.0 was used to call IgBLAST, format the alignments into the Adaptive Immune Receptor Repertoire (AIRR) format, and cluster sequences into clonal groups.^93^ Cells with multiple heavy chain sequences or with only light chain sequences were removed. Clonal abundance/diversity and V-gene usage were calculated with Alakazam v.1.3.0, and SMH rates were calculated with SHazaM v.1.2.0. Lineage trees were reconstructed with the Dowser v2.3 package. All computational analyses were performed in R (version 4.3.2).

### Bulk RNA sequencing, genotyping, and demultiplexing of individual donor samples

Bulk RNA sequencing (Bulk RNA-seq) was performed on total RNA extracted from thawed human PBMCs using TRIzol (ThermoFisher Scientific). Libraries were prepared using the NEBNext Ultra II RNA Library Prep Kit with the Poly(A) mRNA Magnetic Isolation Module, following the manufacturer’s protocol. Sequencing was conducted on the Illumina NextSeq 2000 platform using 100-base single-end reads with 8-base dual indexing. Data were analyzed using the nf-core/rnaseq pipeline (version 1.4.2), implemented on Nextflow (version 20.07.1).

All RNA-seq data were processed using the human reference genome (Ensembl GRCh38). After adapter trimming, ribosomal RNA removal, and duplicate read marking, donor-specific alignments were used for variant calling following the GATK Best Practices workflow^94^ (https://gatk.broadinstitute.org/hc/en-us/articles/360035531192-RNAseq-short-variant-discovery-SNPs-Indels-) for single nucleotide polymorphism (SNP) and indel discovery. To improve base-calling accuracy, base quality score recalibration (BQSR) was performed using read alignments and a publicly available list of common human variants (“known sites”: 36.6 million SNPs with MAF > 0.0005 from the 1000 Genomes Project: [https://sourceforge.net/projects/cellsnp/files/SNPlist/]).^95^ The SNPs were filtered across individual donor samples, and a merged VCF file was generated to represent donor genotypes.

The “cellranger reanalyze” pipeline was used in obtaining the individual donor GEX data by splitting the pooled scRNA-seq GEX data (feature barcode matrix) based on the individual donor barcodes (from vireo). The V(D)J data for individual donors were obtained in a two-step process: first donor fastq files were obtained using 10x Genomics developed “bamtofastq” tool by splitting the alignments (BAM file for pooled gene expression) data and the individual donor barcodes (from vireo). The resulting individual donor fastq files were run through the “cellranger vdj” pipeline to obtain the individual donor B-cell data.

### Monoclonal antibody (mAb) production

Ig heavy and light chain sequences of selected HCV E1E2-specific B cells, obtained via single-cell V(D)J profiling, were synthesized by GeneArt gene synthesis (ThermoFisher Scientific). These synthesized sequences were subsequently cloned into human Igγ1, Igκ and Igλ expression vectors as prviously described.^79^ Plasmids containing paired antibody heavy- and light-chain sequences were co-transfected (1:1 ratio) with pAdvantage (Promega) into Expi293 suspension cells using ExpiFectamine™ 293 Transfection Kit (ThermoFisher Scientific) according to the manufacturer’s protocol. Antibody-containing supernatants were harvested 14 days post transfection by centrifugation to pellet cells, filtered through 0.22 µm filters and purified over Protein A-Sepharose 4 Fast Flow (Cytiva) column according to the manufacturer’s protocol. Purified antibodies were verified by SDS PAGE for purity.

### ELISA

The methods for measuring binding of HCV mAbs to antigens have been described elsewhere.^96^ Briefly, Costar High Binding Half-Area 96-well plates (Corning) were coated overnight at 4 °C with 5 μg/ml of Galanthus nivalis lectin (GNL, Vector Laboratories). After blocking with 4% nonfat milk (Bio-Rad) in PBS + 0.05% Tween-20 (PBS-T), plates were incubated with batch diluted cell lysates from 293T transfected cells expressing E1E2 at room temperature for 1 hour rocking. Serial dilutions mAbs were then added and incubated at room temperature for at least 1 hour rocking. Horseradish peroxidase (HRP)-conjugated goat anti-human IgG Fc fragment antibody (1:2,000, Jackson ImmunoResearch) was used for detection as secondary antibody incubated at room temperature for 1 hour rocking and TMB substrate (ThermoFisher) was used to detect HRP activity that was recorded using a VersaMax microplate reader (Molecular devices) at 450nm absorbance.

For antibody competition analysis, GNL-captured H77 E1E2 in cell lysates from 293T transfected cells was blocked with HCV mAb antibodies (20 µg/mL) and incubated for 30 minutes at room temperature rocking, followed by addition of biotinylated reference mAbs diluted to 75% maximal binding level. Binding was detected using HRP–conjugated streptavidin (1:2,000; Thermo Fisher Scientific). Results are expressed as the percentage of inhibition without blocking antibody.

### HCV neutralization assay

HCV pseudoparticles (HCVpp) were generated by co-transfection of 293T CD81KO cells^97^ with pNL4-3.lucR^-^E^-^, pAdvantage (Promega) and the corresponding expression plasmids encoding the E1E2 from isolate H77 (Tier 2, genotype 1a), 5.2.1 (Tier 1, genotype 5a), 1a123 (Tier 2, genotype 1a), 4.2.2 (Tier 3, genotype 4a) or 3.1.2 (Tier 4, genotype 3a) at a 4:1 ratio by polyethylenimine (PEI, Kyfora Bio). HCVpp neutralization was carried out on Huh-7 cells with mAbs at a desired concentration as previously described.^98^ Virus infectivity was detected with Bright-Glo luciferase assay system (Promega) according to the manufacturer’s protocol and luminescence activity (RLU) was recorded using SpectraMax iD3 (Molecular Devices), and percent neutralization was calculated as the virus infectivity inhibited at the antibody concentrations divided by the infectivity without antibody after background subtraction. The background infectivity of the pseudoparticle virus was defined by infecting cells with virus made only with pNL4-3.lucR-E-. Pseudoparticles displaying the vesicular stomatitis virus envelope glycoprotein G (VSVpp) were used as control for nonspecific neutralizing activity.

### Statistical analysis

Statistical analyses were performed using GraphPad Prism (version 10.2.3), and details are provided in the figure legends. Comparisons between two groups were done using two-tailed Mann-Whitney U-tests and Wilcoxon t-tests for unpaired and paired data, respectively. For each analysis, statistical significance was established at *p*-value < 0.05.

## RESOURCE AVAILABILITY

### Lead contact

Further information and requests for resources and reagents should be directed to and will be fulfilled by the lead contact, Mansun Law.

### Materials availability

Antibodies generated in this study are available from the lead contact with a completed Material Transfer Agreement.

### Data and code availability

All scRNA-seq and bulk RNA-seq data generated in this study have been deposited to Gene Expression Omnibus (GEO) under superseries GES295931.

This study did not generate any original code.

Any additional information required to reanalyze the data reported in this paper is available from the lead contact upon request.

## ACKNOWLEDGEMENT

We thank Hyun Min Kang (University of Michigan) for advice on single-cell data demultiplexing, Sorin Draghici (Advaita Bioinformatics) for advice on iPathwayGuide analysis, Yuxing Li (University of Maryland) for providing the expression plasmid, Natalie Shoanga (Scripps Research) for technical support, and Miyo Ota, David Nemazee and Stefan Wieland (Scripps Research) for discussion. Some clinical samples used in this study were collected in the clinical trial NCT02476617 sponsored by Abbvie. This work was supported in part by the Division of Intramural Research of the National Library of Medicine (NLM), National Institutes of Health (NIH). We acknowledge the support of NIH grants AI158193, AI168251, AI168917, and AI184931 (M.L.), AI086230 (A.Y.K., G.M.L.), AI132213 and AI168048 (T.R.F., E.A.T.). The funding bodies had no role in the design or conclusions of this study. The graphical abstract was created using Biorender.com.

## AUTHOR CONTRIBUTIONS

F.C. and M.L. designed and conceived the study; F.C. conducted flow cytometry analysis and cell sorting; T.M. and S.R.H. performed scRNA-seq, bulk RNA-seq and library preparation; F.C., P.N., and A.S. performed scRNA-seq bioinformatics analysis; L.K., E.A.T., and T.R.F. generated the HCV E1E2 antigen probe; R.A. generated the SARS-CoV-2 spike antigen probe; E.G. and S.C.L. performed mAb characterization; Y.Z. and R.H.S. assisted with single-cell gene expression data analysis; X.L. assisted with Ig sequence analysis; G.M.L., J.A., A.Y.K., L.L.X., and P.P. coordinated clinical cohorts and obtained blood samples; M.L., E.A.T., and T.R.F. provided funding support; F.C. drafted the figures and manuscript. All authors read, edited, and approved the final manuscript.

---

**Figure S1.**
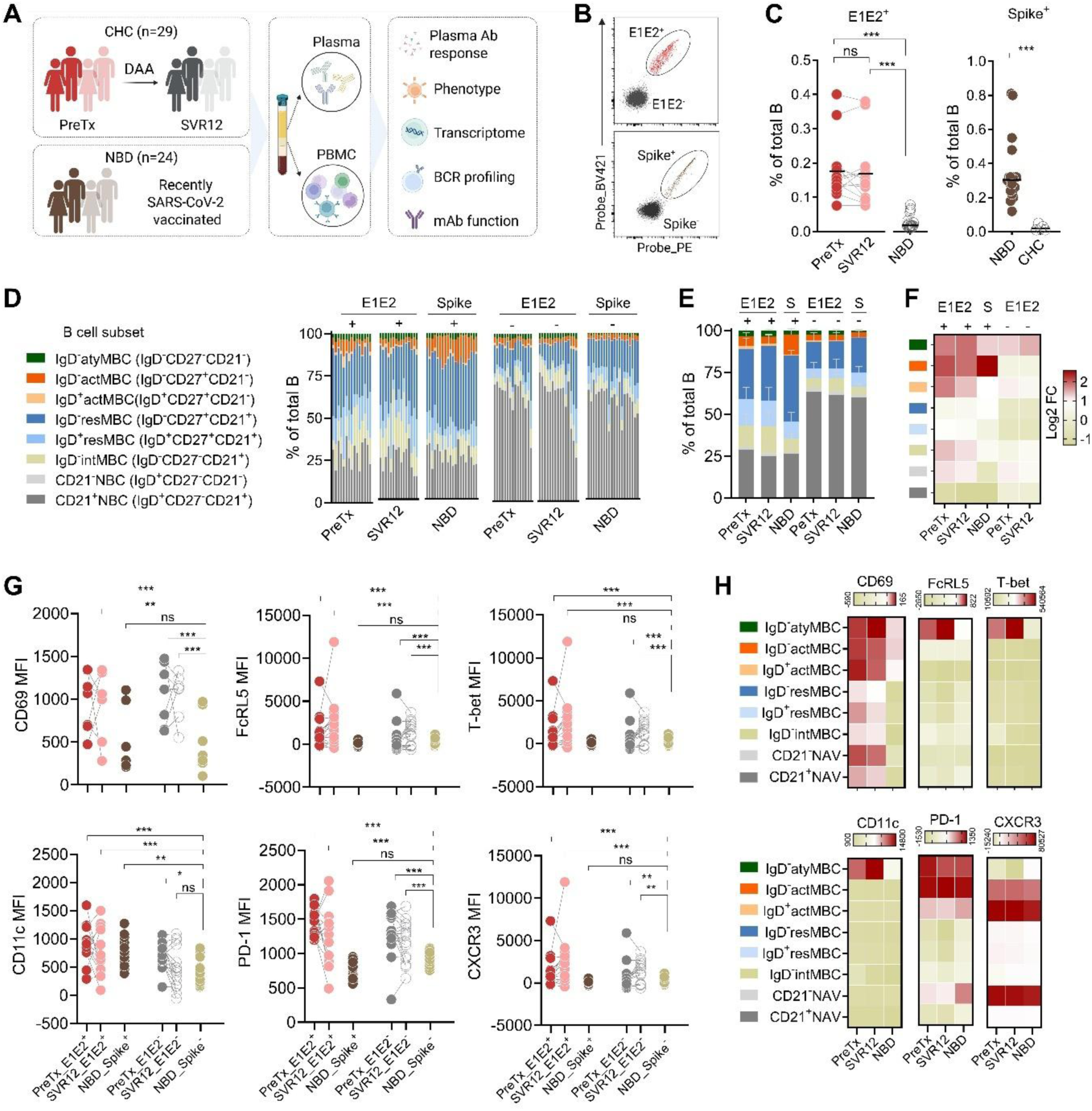
Phenotypic characterization of antigen-specific and non-specific B cells from CHC and NBDs. (A) Schematic of sample cohort and study design. (B) Representative flow cytometry plots for detection of antigen-specific B cells in CHC patients and NBDs. (C) Frequency of antigen-specific B cells in CHC patients and NBDs. (D-E) Distribution of B cell subsets in each cohort, shown as individual (D) and average (E). S, spike. (F) Log2 fold change (FC) of B cell subsets in each group relative to healthy control level (spike^-^ B cells from NBDs). (G) Mean fluorescence intensity (MFI) quantifying the expression of activation and dysregulation markers in each group. (H) Heatmap detailing the expression level of each marker across different B cell subsets. *P* values were determined by Wilcoxon’s paired t test (A) and tow-tailed Mann-Whitney test (F) and. *P < 0.05, ** P < 0.01, ***P < 0.001, ns, not significant.

**Figure S2.**
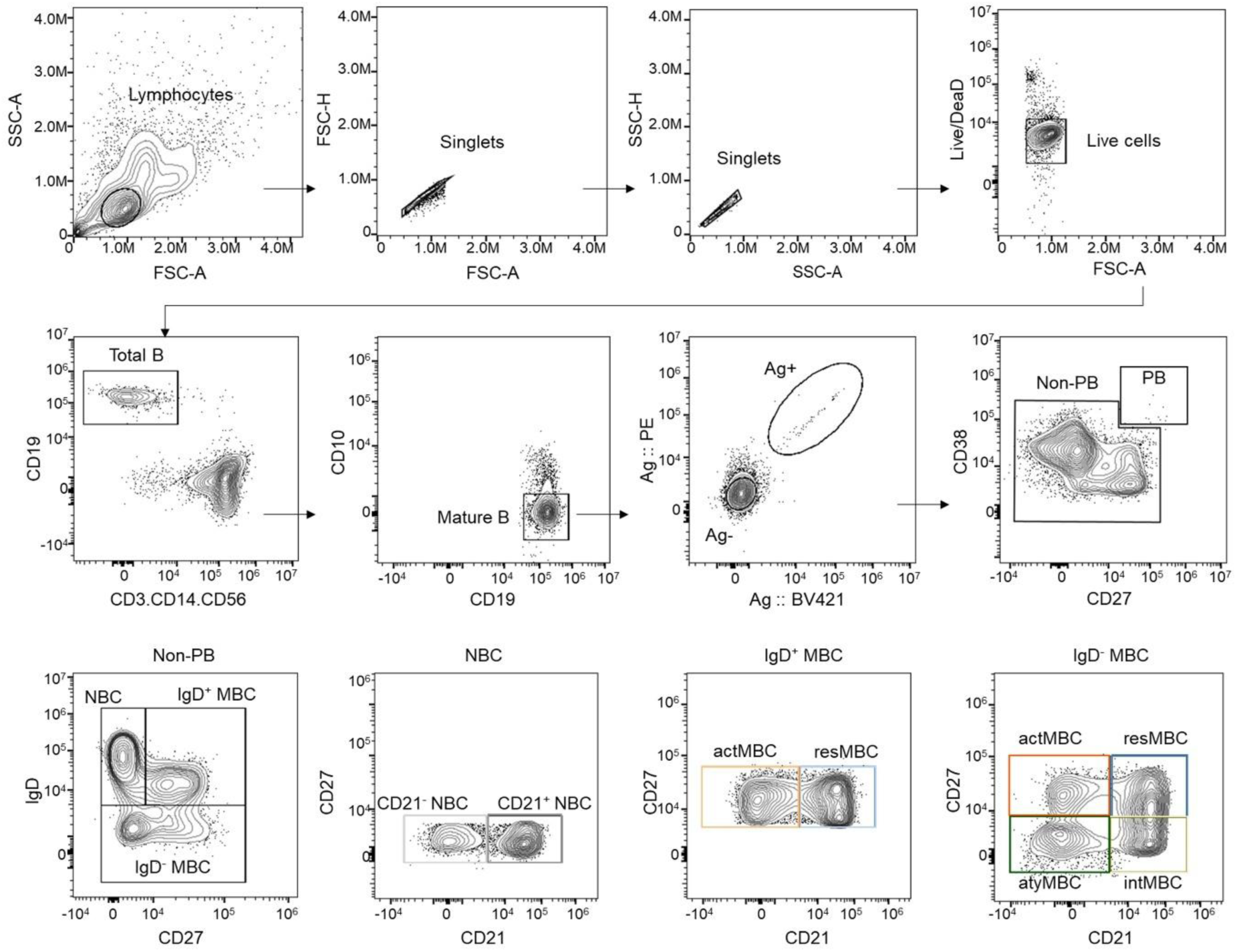
Flow cytometry gating strategy for analysis of antigen-specific and non-specific B cells in CHC and NBDs, related to Figure S1. Ag, antigen; PB, plasmablasts.

**Figure S3.**
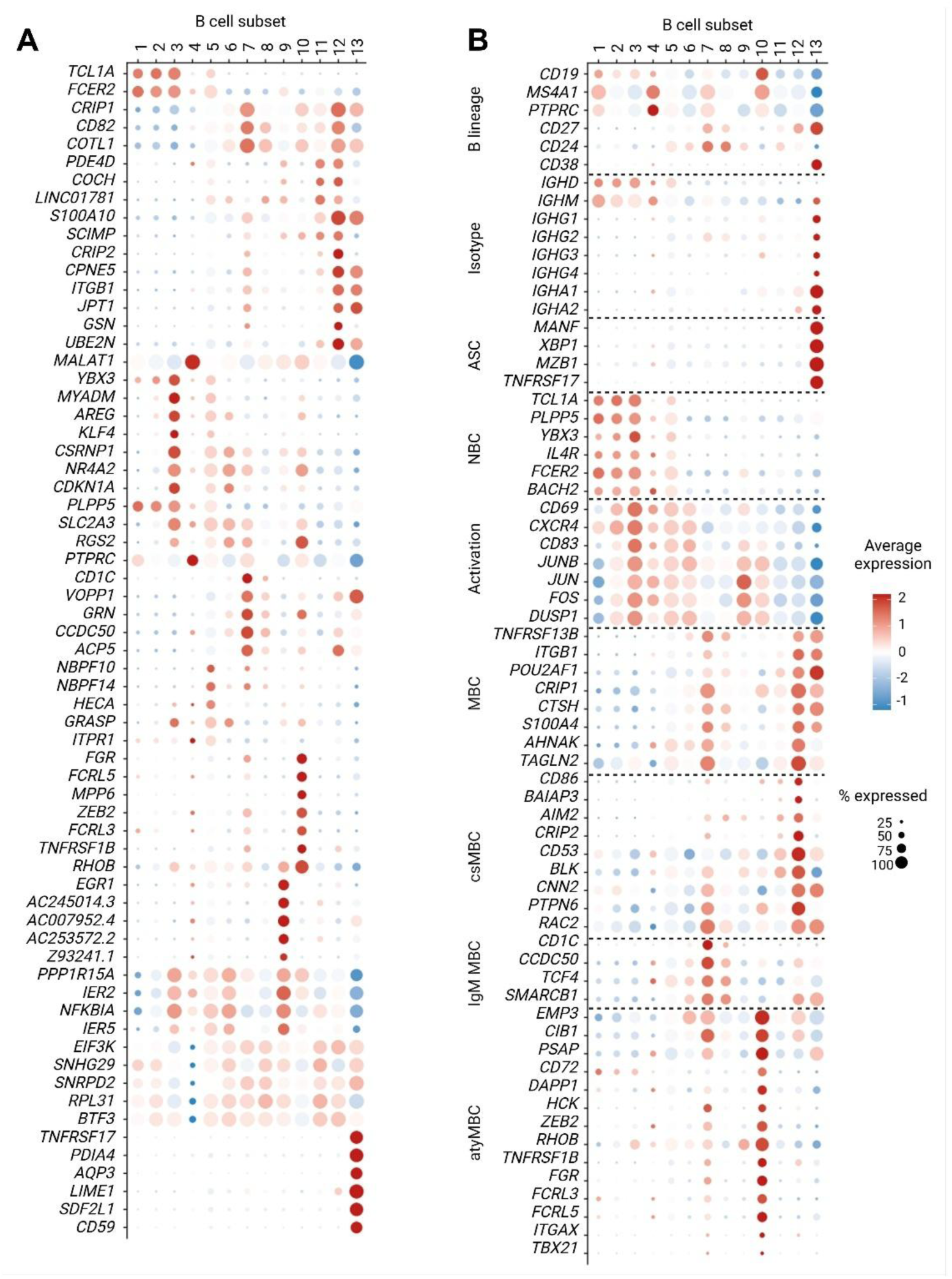
Identification of B cell subsets by single-cell transcriptional analysis, related to. **Figure 1A**. (A) Dot plots showing differentially expressed genes (DEGs) in each subset. (B) Annotation of B cell subsets using previously documented B cell transcriptional profiles.

**Figure S4.**
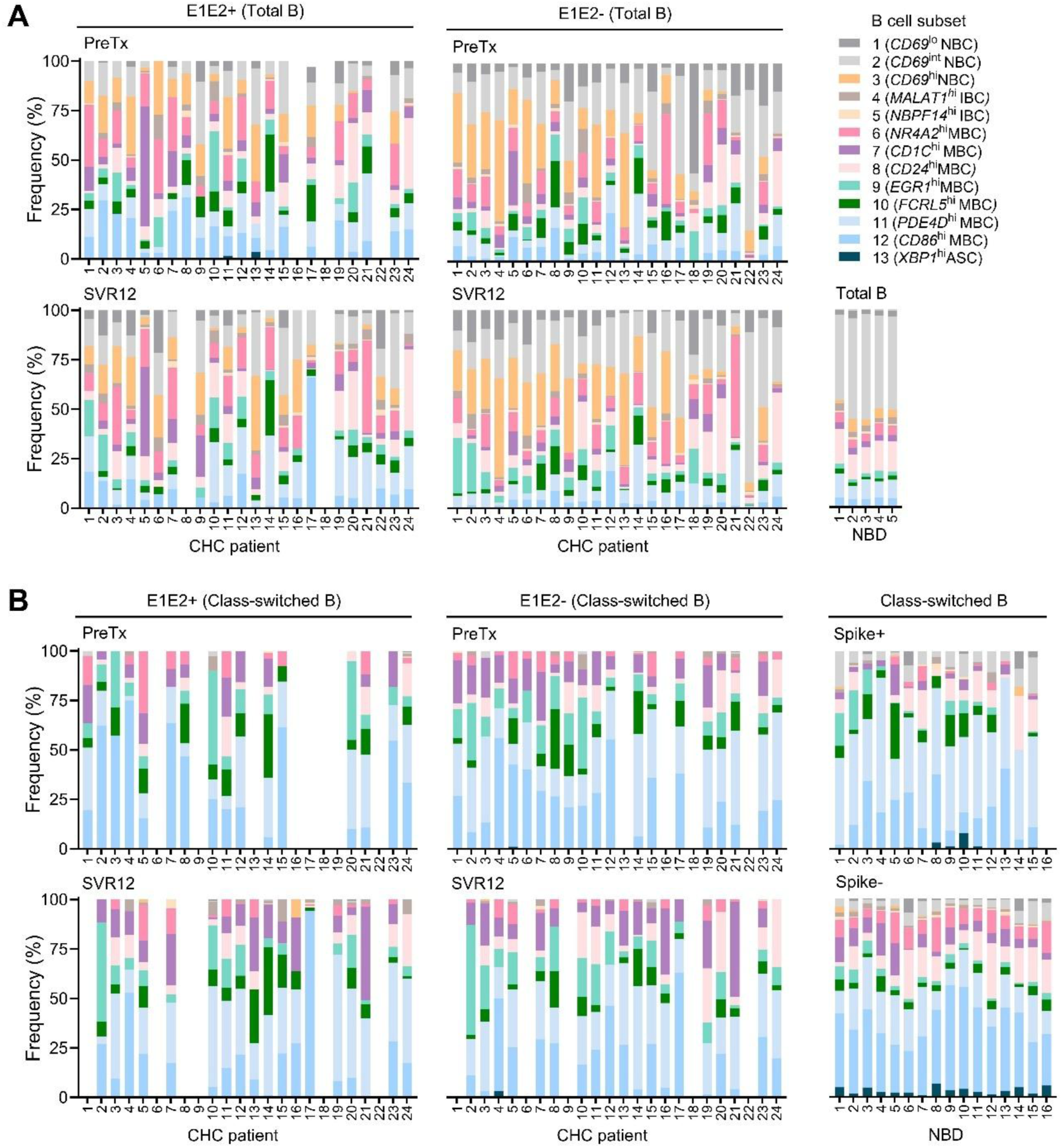
B cell cluster distribution in individual CHC and NBDs, related to Figure 1F. (A) Distribution of subsets among total B cells. (B) Distribution of subsets among class-switched B cells. To minimize bias, data from individual subjects with total sequence counts fewer than 50 were excluded.

**Figure S5.**
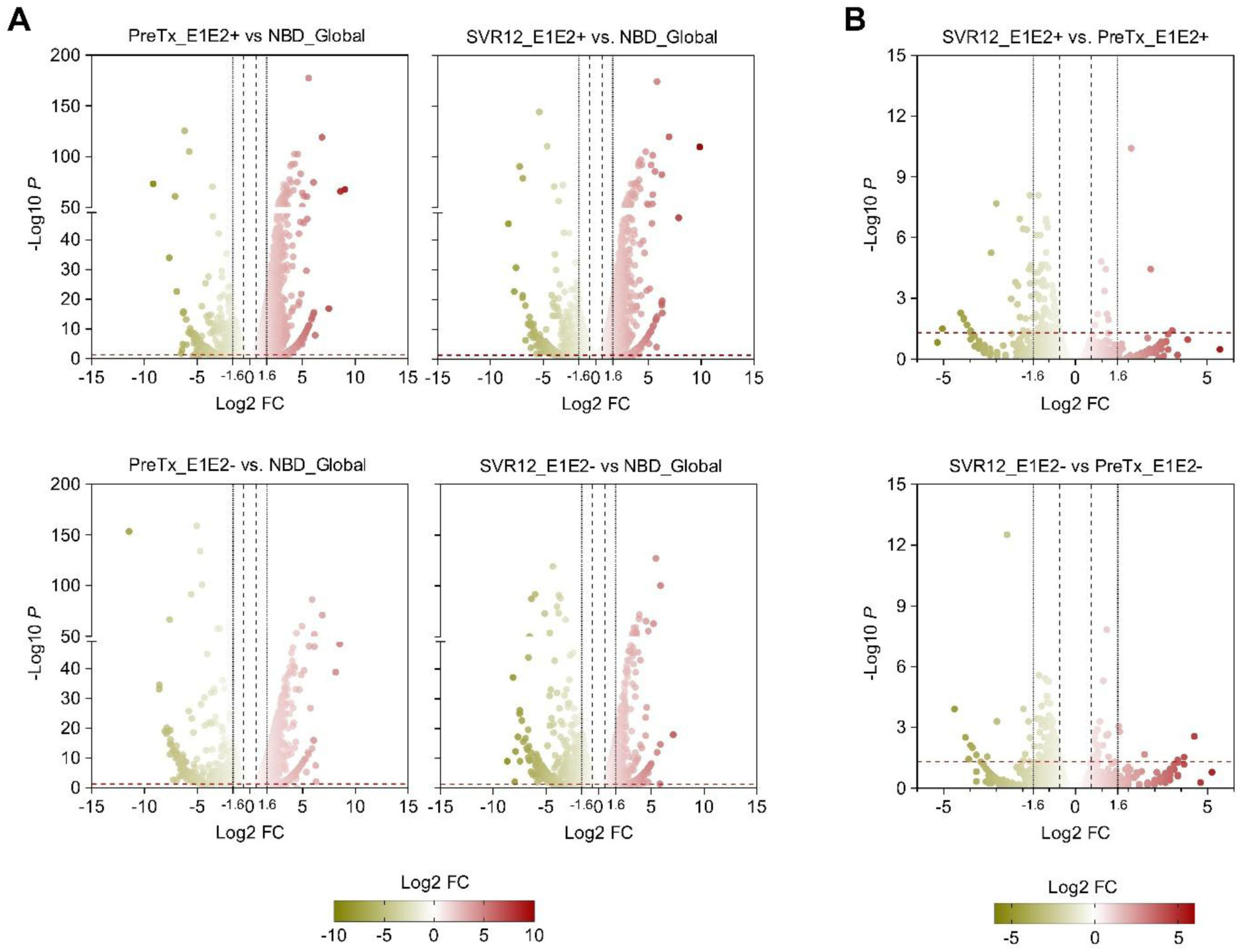
DEG profiles in B cells from CHC across comparisons. Related to Figure 3. Volcano plots showing all genes with measured expression in the indicated comparisons. Red horizontal dashed lines indicate the significance cutoff (–log₁₀(*p*) = 1.3; *p* = 0.05), and black vertical dashed lines indicate log₂ FC thresholds of |0.6| and |1.6| (fold change = |1.5| and |3|). FC, fold change.

**Figure S6.**
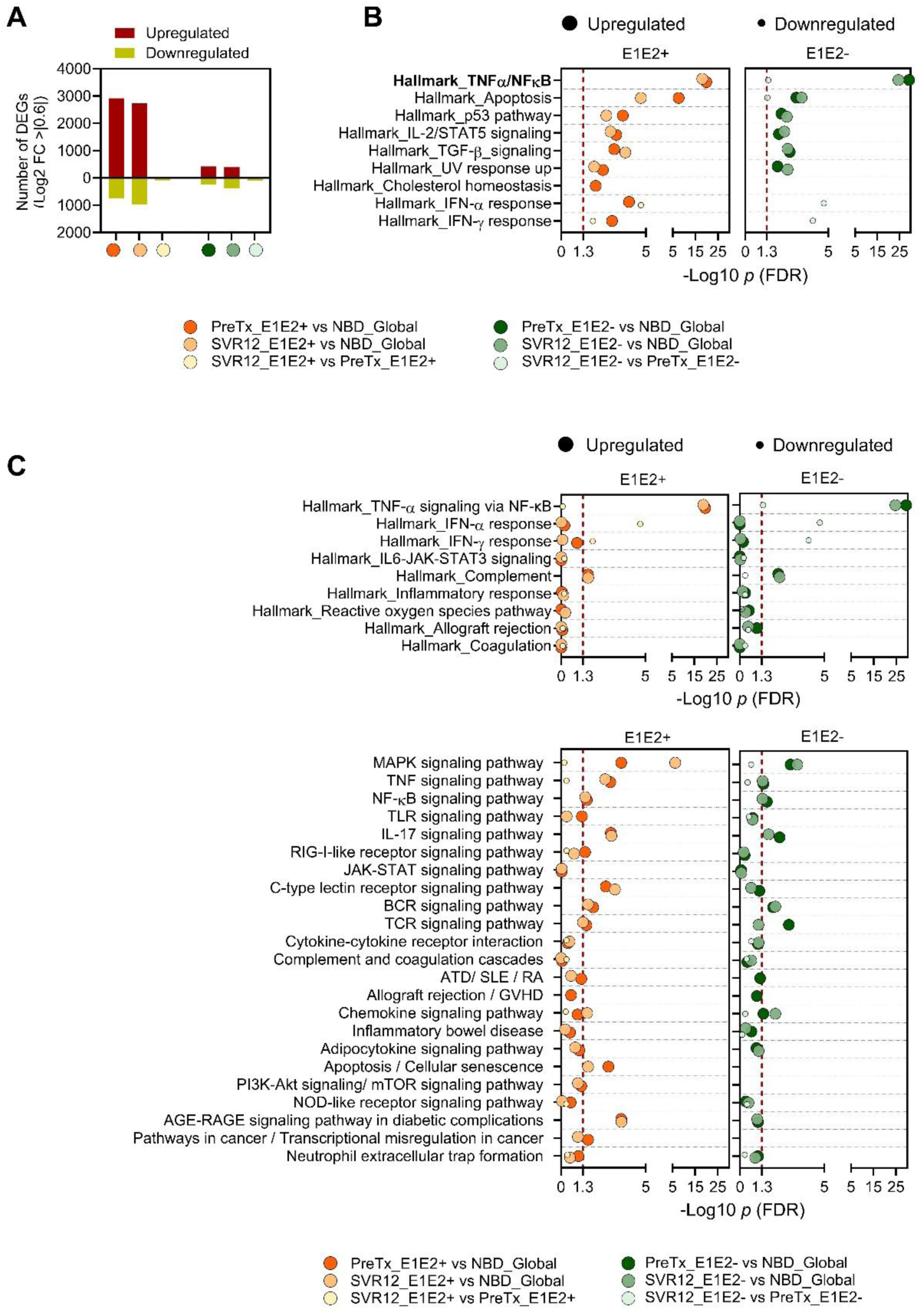
Gene set and pathway analysis of DEGs in CHC B cells. Related to Figure 3. (A) Number of DEGs identified in each comparison. DEG thresholds: |log₂FC|> 0.6 (fold change >1.5) and –log₁₀(p-value) >1.3 (p-value < 0.05). (B) Significantly altered hallmark signatures (MSigDB) in each comparison, with statistical significance assessed using false discovery rate (FDR)-adjusted *p*-values. Doted lines indicate the -Log10 (*p*-value) threshold of 1.3, corresponding to a *p*-value of 0.05. (C) Altered pathways associated with chronic inflammation in CHC before and after treatment. Upper panel, Inflammation-related hallmark signatures from MSigDB. Botten panel, Inflammation-related pathways from KEGG. Statistical significance was determined using FDR-adjusted *p*-values. Dotted lines indicate -Log₁₀ (*p*-value) threshold of 1.3, corresponding to a *p*-value of 0.05.

**Figure S7.**
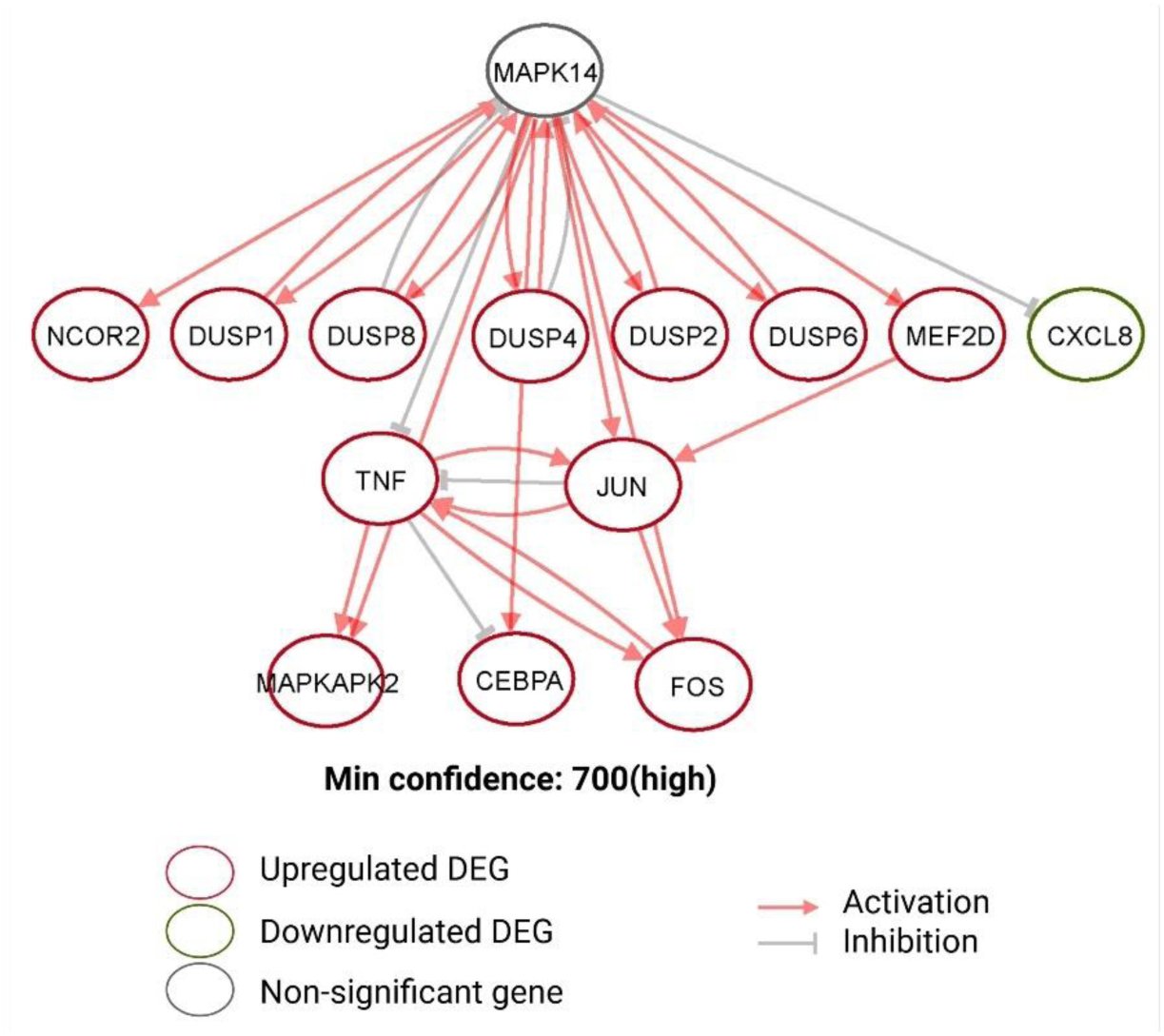
MAPK14 serves as a potential upstream regulator of DEGs in E1E2^+^ B cells from CHC compared to NBDs, related to Figure 3F.

**Figure S8.**
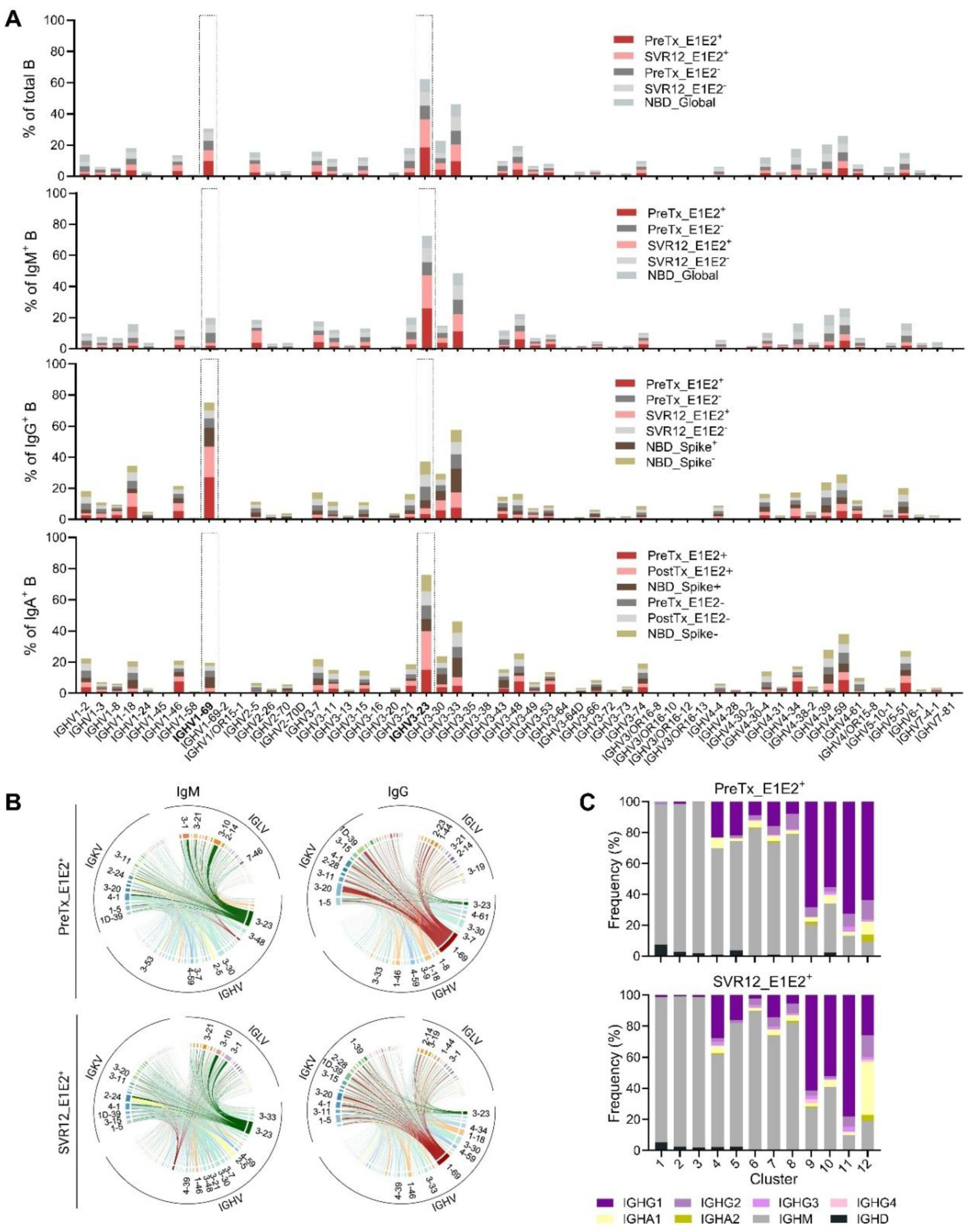
Genetic features of BCR repertoire in CHC and NBDs, related to Figure 5A. (A) IGHV gene usage in total, IgM-, IgG- and IgA-expressing B cells from CHC and NBDs. (B) Paired heavy and light chain V gene usage in E1E2⁺ IgM and IgG B cells from CHC before and after treatment. (C) Isotype distribution across E1E2⁺ B cell subsets from CHC before and after treatment.

**Figure S9.**
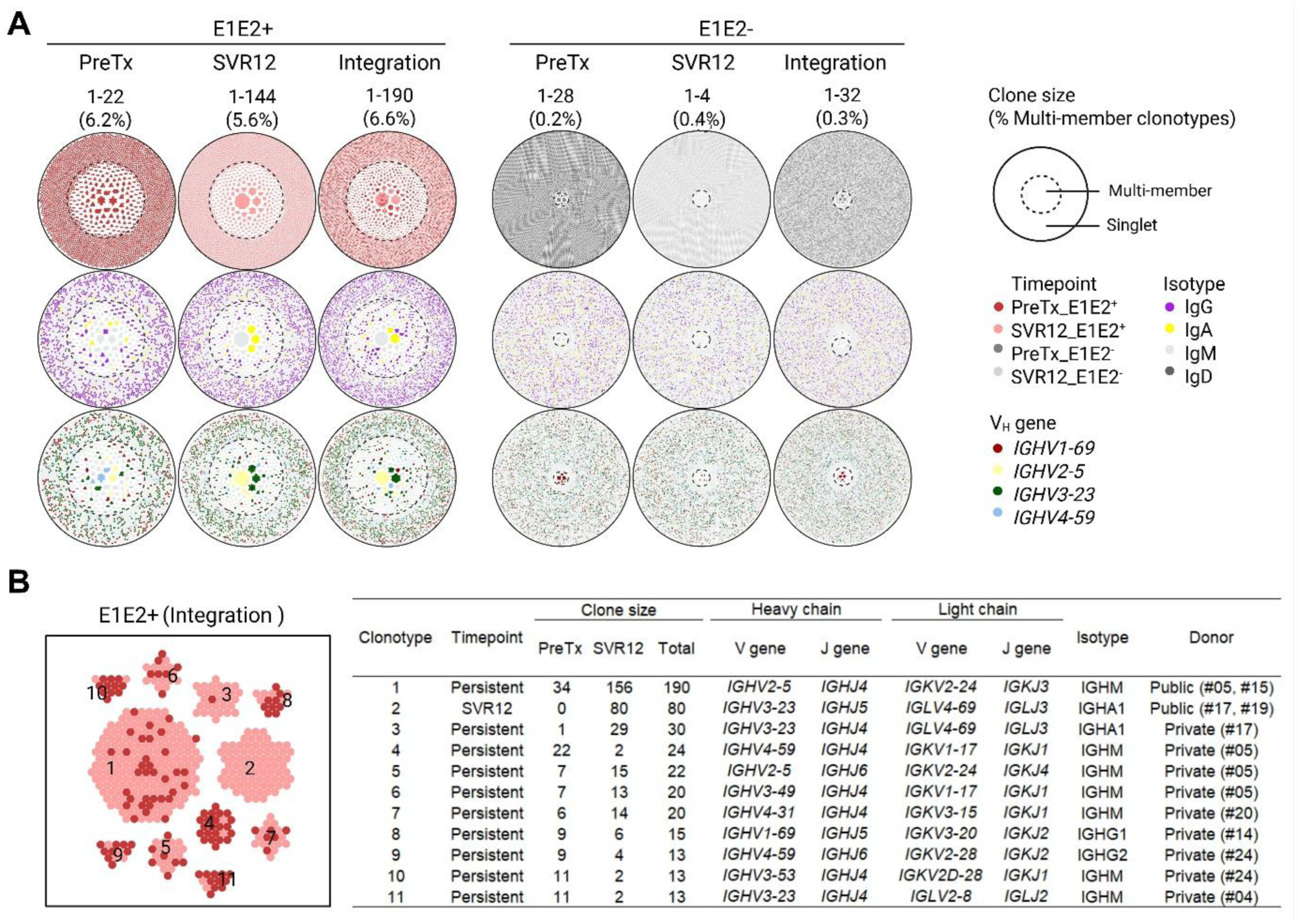
Clonotypes in the BCR repertoire of CHC before and after treatment, related to Figure 5. (A) Clonotypes and their corresponding isotype and IGHV gene usage in E1E2^+^ and E1E2^-^ B cells from CHC. (B) General information of the top 11 clonal families in E1E2^+^ B cells from CHC.

**Figure S10.**
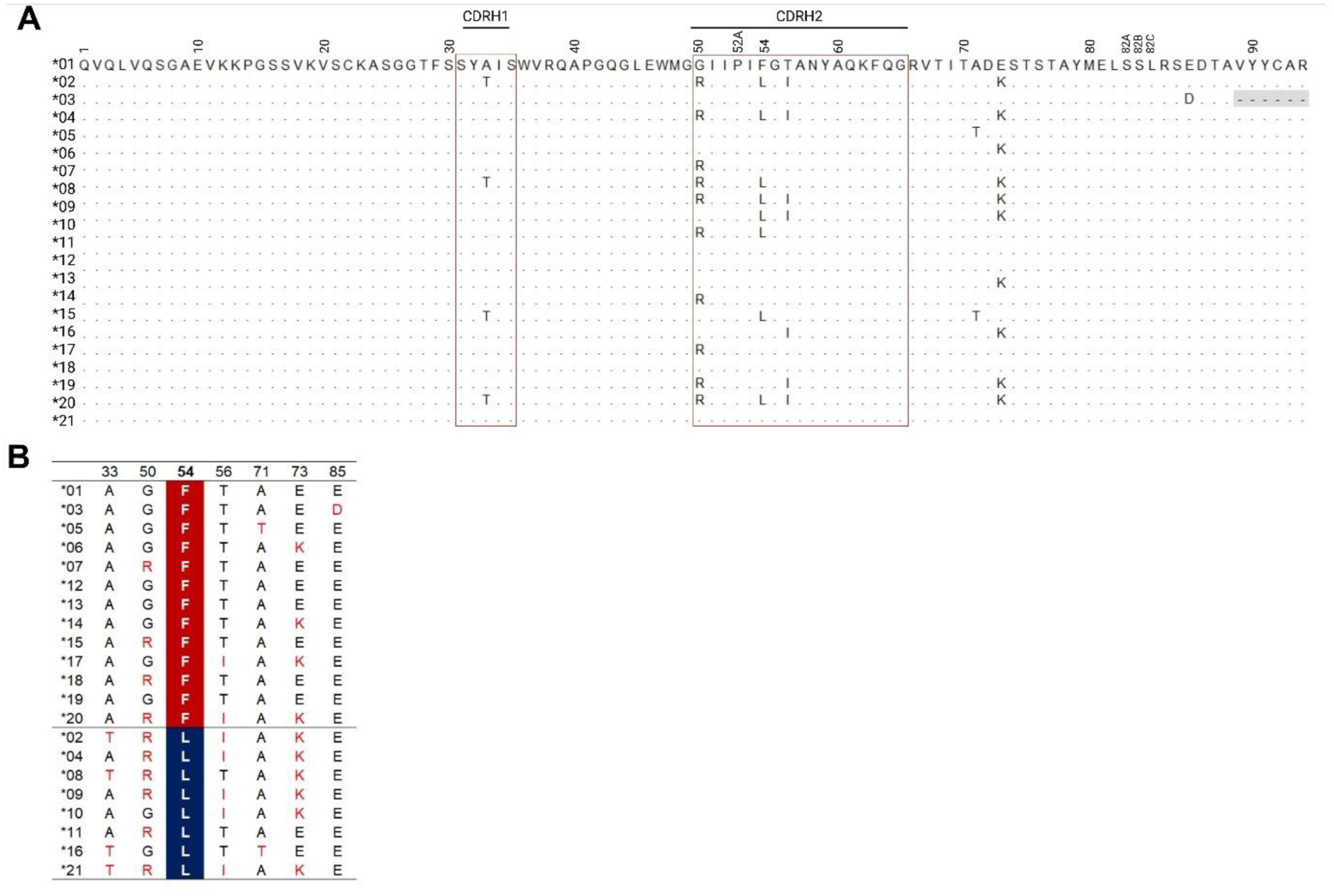
Human *IGHV1-69* gene alleles, related to Figures 6D and 6H. (A) Amino acid sequence alignment of *IGHV1-69* alleles. (B) Summary of polymorphic amino acid positions across *IGHV1-69* alleles.

**Figure S11.**
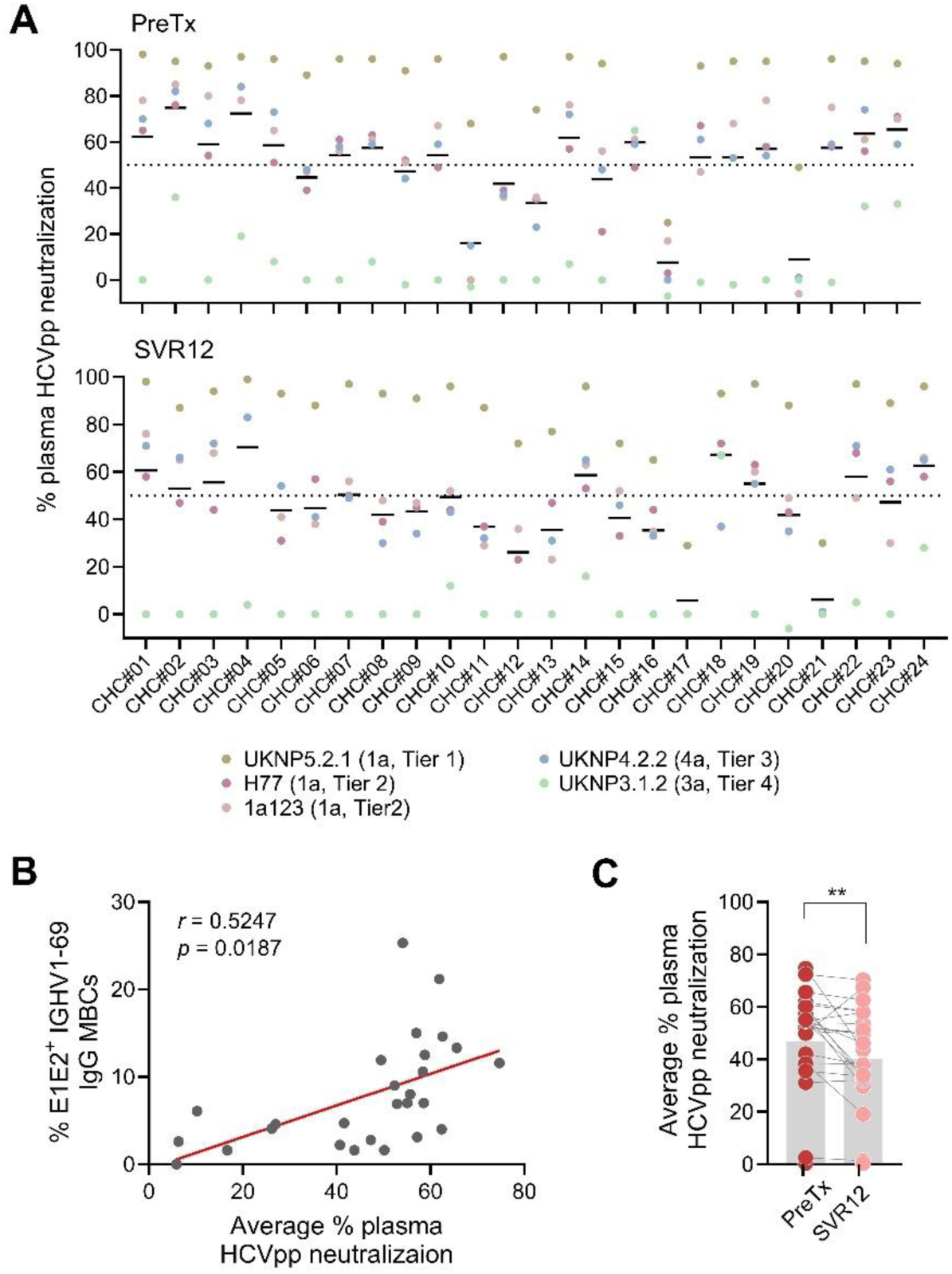
Plasma neutralizing activity in CHC individuals before and after treatment, related to Figure 6E. (A) Plasma neutralization against a panel of 5 selected HCV pseudoparticles (HCVpp) spanning 4 tiers of neutralization resistance. (B) Linear regression analysis of frequency of E1E2^+^ total B cells and the average plasma neutralization in CHC patients. (C) Dynamics of average plasma neutralization (% HCVpp infection inhibition) in CHC individuals before and after treatment. Each dot represents a single donor. Statistical significance was determined using paired Wilcoxon t-tests. **, *p*-value <0.01.

**Table S1.** Clinical parameters of CHC patients undergoing DAA treatment.

| Patient ID | Gender | Age | Exposure years | HCV Genotype | Viral load (IU/mL) |  | Complications | scRNA-seq | Flow cytometry |
| --- | --- | --- | --- | --- | --- | --- | --- | --- | --- |
|  |  |  |  |  | PreTx | SVR12 |  |  |  |
| CHC#01 | Male | 42 | 30 | 1a | 1,230,000 | Yes | Non-Hodgkins lymphoma | √ | √ |
| CHC#02 | Male | 49 | 9 | 1a | 6,360,000 | Yes | Nothing significant | √ | √ |
| CHC#03 | Male | 32 | 10 | 1a | 75,600 | Yes | Hepatic lesion | √ | √ |
| CHC#04 | Male | 57 | 37 | 1a | 40,000 | Yes | Nothing significant | √ | √ |
| CHC#05 | Male | 44 | 14 | 1a | 4,270,000 | Yes | Nothing significant | √ | √ |
| CHC#06 | Male | 38 | 12 | 1a | 1,320,000 | Yes | Fatty liver | √ | √ |
| CHC#07 | Female | 51 | 16 | 1a | 4,570,000 | Yes | Nothing significant | √ | √ |
| CHC#08 | Male | 64 | 5 | 1a | 300,000 | Yes | Cirrhosis | √ | √ |
| CHC#09 | Female | 72 | 34 | 1a | 186,000 | Yes | Nothing significant | √ | √ |
| CHC#10 | Male | 29 | 3 | 1a | 1,360,000 | Yes | Nothing significant | √ | √ |
| CHC#11 | Male | 29 | > 1 | 1a | 258,000 | Yes | Nothing significant | √ |  |
| CHC#12 | Male | 42 | 5 | 1a | 597,000 | Yes | Nothing significant | √ |  |
| CHC#13 | Female | 63 | 5 | 1a | 1,320,000 | Yes | Nothing significant | √ |  |
| CHC#14 | Female | 61 | 43 | 2 | 2,020,000 | Yes | Nothing significant | √ |  |
| CHC#15 | Female | 45 | 4 | 2b | 1,310,000 | Yes | Type 2 diabetes mellitus with hyperglycemia | √ |  |
| CHC#16 | Male | 59 | 39 | 2b | 1,550,000 | Yes | Nothing significant | √ |  |
| CHC#17 | Female | 63 | > 20 | 1a | 7,350,000 | Yes | Hepatocellular carcinoma | √ |  |
| CHC#18 | Female | 32 | 2 | 1a | 43,700 | Yes | Nothing significant | √ |  |
| CHC#19 | Female | 63 | > 6 | 1a | 3,680,000 | Yes | Nothing significant | √ |  |
| CHC#20 | Male | 46 | 2 | 1a | 1,270,000 | Yes | Nothing significant | √ |  |
| CHC#21 | Male | 56 | 39 | 1b | 1,270,000 | Yes | Cirrhosis | √ |  |
| CHC#22 | Male | 60 | 4 | 1a | 1,350,000 | Yes | Nothing significant | √ |  |
| CHC#23 | Female | 38 | 38 | 1a | 401,000 | Yes | Nothing significant | √ |  |
| CHC#24 | Male | 36 | 10 | 1a | 828,000 | Yes | Nothing significant | √ |  |
| CHC#25 | M | 65 | 20 | 1b | 2,510,000 | Yes | HCC |  | √ |
| CHC#26 | M | 44 | 26 | 1a | 261,000 | Yes | Cirrhosis of liver |  | √ |
| CHC#27 | M | 46 | 28 | 1b | 2,040,000 | Yes | Cirrhosis |  | √ |
| CHC#28 | M | 43 | 6 | 3 | 1,040,000 | Yes | Renal cell cancer |  | √ |
| CHC#29 | F | 45 | 24 | 1b | 5,070,000 | Yes | Fatty liver |  | √ |

**Table S2.**
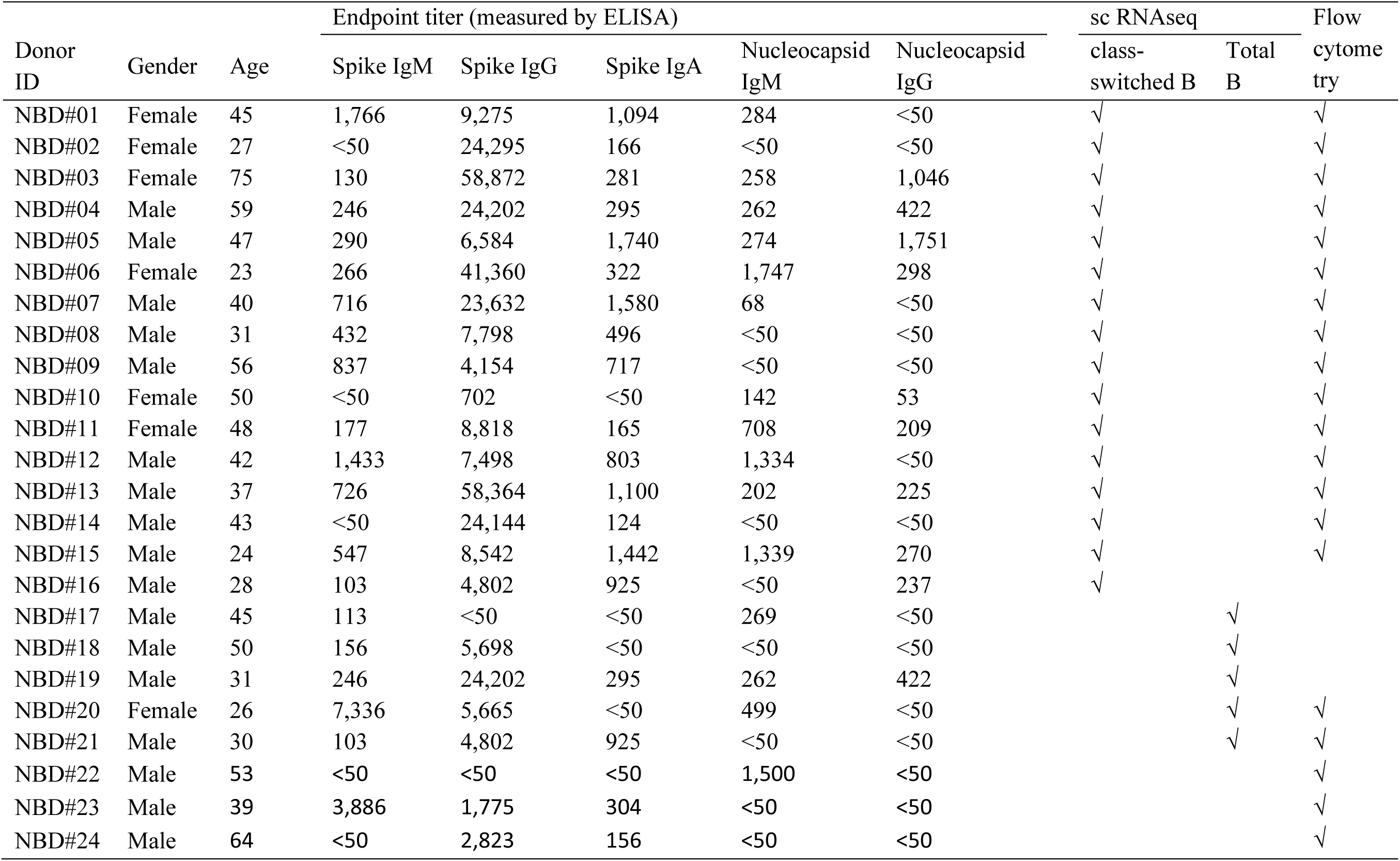
Plasma antibody titers against SARS-CoV2 spike and nucleocapsid proteins in vaccinated NBDs.

**Table S3.**
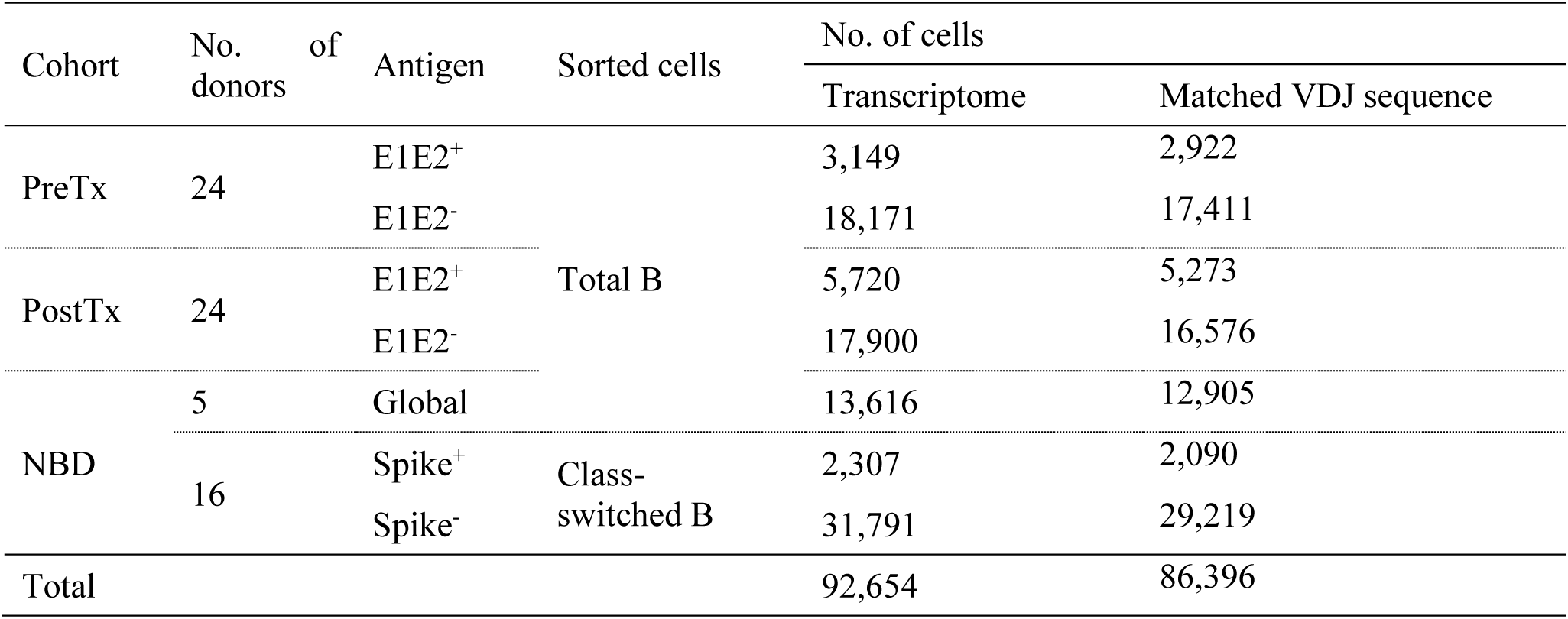
Summary of B cells with matched transcriptome and V(D)J sequences from scRNA-seq.

## REFERENCES

1. Akkaya, M., Kwak, K., and Pierce, S.K. (2020). B cell memory: building two walls of protection against pathogens. Nat. Rev. Immunol. 20, 229–238. 10.1038/s41577-019-0244-2.

2. Upasani, V., Rodenhuis-Zybert, I., and Cantaert, T. (2021). Antibody-independent functions of B cells during viral infections. PLoS Pathog. 17, e1009708. 10.1371/journal.ppat.1009708.

3. Nothelfer, K., Sansonetti, P.J., and Phalipon, A. (2015). Pathogen manipulation of B cells: the best defence is a good offence. Nat. Rev. Microbiol. 13, 173–184. 10.1038/nrmicro3415.

4. Cooper, L., and Good-Jacobson, K.L. (2020). Dysregulation of humoral immunity in chronic infection. Immunol. Cell Biol. 98, 456–466. 10.1111/imcb.12338.

5. Martinello, M., Solomon, S.S., Terrault, N.A., and Dore, G.J. (2023). Hepatitis C. Lancet 402, 1085–1096. 10.1016/S0140-6736(23)01320-X.

6. Law, M. (2021). Antibody responses in hepatitis C infection. Cold Spring Harb. Perspect. Med. 11. 10.1101/cshperspect.a036962.

7. Salinas, E., Boisvert, M., Upadhyay, A.A., Bedard, N., Nelson, S.A., Bruneau, J., Derdeyn, C.A., Marcotrigiano, J., Evans, M.J., Bosinger, S.E., et al. (2021). Early T follicular helper cell activity accelerates hepatitis C virus-specific B cell expansion. J. Clin. Invest. 131. 10.1172/JCI140590.

8. Nishio, A., Hasan, S., Park, H., Park, N., Salas, J.H., Salinas, E., Kardava, L., Juneau, P., Frumento, N., Massaccesi, G., et al. (2022). Serum neutralization activity declines but memory B cells persist after cure of chronic hepatitis C. Nat Commun 13, 5446. 10.1038/s41467-022-33035-z.

9. Ogega, C.O., Skinner, N.E., Flyak, A.I., Clark, K.E., Board, N.L., Bjorkman, P.J., Crowe, J.E., Jr., Cox, A.L., Ray, S.C., and Bailey, J.R. (2022). B cell overexpression of FCRL5 and PD-1 is associated with low antibody titers in HCV infection. PLoS Pathog. 18, e1010179. 10.1371/journal.ppat.1010179.

10. Ambegaonkar, A.A., Holla, P., Dizon, B.L., Sohn, H., and Pierce, S.K. (2022). Atypical B cells in chronic infectious diseases and systemic autoimmunity: Puzzles with many missing pieces. Curr. Opin. Immunol. 77, 102227. 10.1016/j.coi.2022.102227.

11. Keller, B., and Warnatz, K. (2023). T-bet^high^CD21^low^ B cells: The need to unify our understanding of a distinct B cell population in health and disease. Curr. Opin. Immunol. 82, 102300. 10.1016/j.coi.2023.102300.

12. Young, C., Singh, M., Jackson, K.J.L., Field, M.A., Peters, T.J., Angioletti-Uberti, S., Frenkel, D., Ravishankar, S., Gupta, M., Wang, J.J., et al. (2025). A triad of somatic mutagenesis converges in self-reactive B cells to cause a virus-induced autoimmune disease. Immunity. 10.1016/j.immuni.2024.12.011.

13. Cornberg, M., Mischke, J., Kraft, A.R., and Wedemeyer, H. (2023). Immunological scars after cure of hepatitis C virus infection: Long-HepC? Curr. Opin. Immunol. 82, 102324. 10.1016/j.coi.2023.102324.

14. Sepulveda-Crespo, D., Volpi, C., Amigot-Sanchez, R., Yelamos, M.B., Diez, C., Gomez, J., Hontanon, V., Berenguer, J., Gonzalez-Garcia, J., Martin-Escolano, R., et al. (2024). Sustained Long-Term Decline in Anti-HCV Neutralizing Antibodies in HIV/HCV-Coinfected Patients Five Years after HCV Therapy: A Retrospective Study. Pharmaceuticals (Basel) 17. 10.3390/ph17091152.

15. Amanna, I.J., Carlson, N.E., and Slifka, M.K. (2007). Duration of humoral immunity to common viral and vaccine antigens. N. Engl. J. Med. 357, 1903–1915. 10.1056/NEJMoa066092.

16. Hajarizadeh, B., Cunningham, E.B., Valerio, H., Martinello, M., Law, M., Janjua, N.Z., Midgard, H., Dalgard, O., Dillon, J., Hickman, M., et al. (2020). Hepatitis C reinfection after successful antiviral treatment among people who inject drugs: A meta-analysis. J. Hepatol. 72, 643–657. 10.1016/j.jhep.2019.11.012.

17. Hosseini-Hooshyar, S., Hajarizadeh, B., Bajis, S., Law, M., Janjua, N.Z., Fierer, D.S., Chromy, D., Rockstroh, J.K., Martin, T.C.S., Ingiliz, P., et al. (2022). Risk of hepatitis C reinfection following successful therapy among people living with HIV: A global systematic review, meta-analysis, and meta-regression. Lancet HIV 9, e414–e427. 10.1016/S2352-3018(22)00077-7.

18. Negro, F. (2021). Residual risk of liver disease after hepatitis C virus eradication. J. Hepatol. 74, 952–963. 10.1016/j.jhep.2020.11.040.

19. Celli, R., Saffo, S., Kamili, S., Wiese, N., Hayden, T., Taddei, T., and Jain, D. (2021). Liver pathologic changes after direct-acting antiviral agent therapy and sustained virologic response in the setting of chronic hepatitis C virus infection. Arch. Pathol. Lab. Med. 145, 419–427. 10.5858/arpa.2020-0008-OA.

20. Cacoub, P., Comarmond, C., Vieira, M., Regnier, P., and Saadoun, D. (2022). HCV-related lymphoproliferative disorders in the direct-acting antiviral era: From mixed cryoglobulinaemia to B-cell lymphoma. J. Hepatol. 76, 174–185. 10.1016/j.jhep.2021.09.023.

21. Cacoub, P., and Saadoun, D. (2021). Extrahepatic Manifestations of Chronic HCV Infection. N. Engl. J. Med. 384, 1038–1052. 10.1056/NEJMra2033539.

22. Courey-Ghaouzi, A.D., Kleberg, L., and Sundling, C. (2022). Alternative B Cell Differentiation During Infection and Inflammation. Front. Immunol. 13, 908034. 10.3389/fimmu.2022.908034.

23. Popescu, M., Cabrera-Martinez, B., and Winslow, G.M. (2019). TNF-α contributes to lymphoid tissue disorganization and germinal center B cell suppression during intracellular bacterial infection. J. Immunol. 203, 2415–2424. 10.4049/jimmunol.1900484.

24. Figgett, W.A., Vincent, F.B., Saulep-Easton, D., and Mackay, F. (2014). Roles of ligands from the TNF superfamily in B cell development, function, and regulation. Semin. Immunol. 26, 191–202. 10.1016/j.smim.2014.06.001.

25. Aaron, T., Laudermilch, E., Benet, Z., Ovando, L.J., Chandran, K., and Fooksman, D. (2023). TNF-α limits serological memory by disrupting the bone marrow niche. J. Immunol. 210, 595–608. 10.4049/jimmunol.2200053.

26. Guest, J.D., Wang, R., Elkholy, K.H., Chagas, A., Chao, K.L., Cleveland, T.E.t., Kim, Y.C., Keck, Z.Y., Marin, A., Yunus, A.S., et al. (2021). Design of a native-like secreted form of the hepatitis C virus E1E2 heterodimer. Proc. Natl. Acad. Sci. U. S. A. 118. 10.1073/pnas.2015149118.

27. Metcalf, M.C., Janus, B.M., Yin, R., Wang, R., Guest, J.D., Pozharski, E., Law, M., Mariuzza, R.A., Toth, E.A., Pierce, B.G., et al. (2023). Structure of engineered hepatitis C virus E1E2 ectodomain in complex with neutralizing antibodies. Nat Commun 14, 3980. 10.1038/s41467-023-39659-z.

28. Tzarum, N., Wilson, I.A., and Law, M. (2018). The neutralizing face of hepatitis C virus E2 envelope glycoprotein. Front. Immunol. 9, 1315. 10.3389/fimmu.2018.01315.

29. Torrents de la Pena, A., Sliepen, K., Eshun-Wilson, L., Newby, M.L., Allen, J.D., Zon, I., Koekkoek, S., Chumbe, A., Crispin, M., Schinkel, J., et al. (2022). Structure of the hepatitis C virus E1E2 glycoprotein complex. Science 378, 263–269. 10.1126/science.abn9884.

30. Giang, E., Dorner, M., Prentoe, J.C., Dreux, M., Evans, M.J., Bukh, J., Rice, C.M., Ploss, A., Burton, D.R., and Law, M. (2012). Human broadly neutralizing antibodies to the envelope glycoprotein complex of hepatitis C virus. Proc. Natl. Acad. Sci. U. S. A. 109, 6205–6210. 10.1073/pnas.1114927109.

31. Boisvert, M., Zhang, W., Elrod, E.J., Bernard, N.F., Villeneuve, J.P., Bruneau, J., Marcotrigiano, J., Shoukry, N.H., and Grakoui, A. (2016). Novel E2 glycoprotein tetramer detects hepatitis C virus-specific memory B cells. J. Immunol. 197, 4848–4858. 10.4049/jimmunol.1600763.

32. Chen, F., Tzarum, N., Wilson, I.A., and Law, M. (2019). V_H_1-69 antiviral broadly neutralizing antibodies: Genetics, structures, and relevance to rational vaccine design. Curr. Opin. Virol. 34, 149–159. 10.1016/j.coviro.2019.02.004.

33. Weber, T., Potthoff, J., Bizu, S., Labuhn, M., Dold, L., Schoofs, T., Horning, M., Ercanoglu, M.S., Kreer, C., Gieselmann, L., et al. (2022). Analysis of antibodies from HCV elite neutralizers identifies genetic determinants of broad neutralization. Immunity 55, 341–354 e347. 10.1016/j.immuni.2021.12.003.

34. Bailey, J.R., Flyak, A.I., Cohen, V.J., Li, H., Wasilewski, L.N., Snider, A.E., Wang, S., Learn, G.H., Kose, N., Loerinc, L., et al. (2017). Broadly neutralizing antibodies with few somatic mutations and hepatitis c virus clearance. JCI Insight 2, e92872. 10.1172/jci.insight.92872.

35. Merat, S.J., Bru, C., van de Berg, D., Molenkamp, R., Tarr, A.W., Koekkoek, S., Kootstra, N.A., Prins, M., Ball, J.K., Bakker, A.Q., et al. (2019). Cross-genotype AR3-specific neutralizing antibodies confer long-term protection in injecting drug users after hcv clearance. J. Hepatol. 71, 14–24. 10.1016/j.jhep.2019.02.013.

36. Stewart, A., Ng, J.C., Wallis, G., Tsioligka, V., Fraternali, F., and Dunn-Walters, D.K. (2021). Single-cell transcriptomic analyses define distinct peripheral B cell subsets and discrete development pathways. Front. Immunol. 12, 602539. 10.3389/fimmu.2021.602539.

37. Brewer, R.C., Ramadoss, N.S., Lahey, L.J., Jahanbani, S., Robinson, W.H., and Lanz, T.V. (2022). BNT162b2 vaccine induces divergent B cell responses to SARS-CoV-2 S1 and S2. Nat. Immunol. 23, 33–39. 10.1038/s41590-021-01088-9.

38. Dugan, H.L., Stamper, C.T., Li, L., Changrob, S., Asby, N.W., Halfmann, P.J., Zheng, N.Y., Huang, M., Shaw, D.G., Cobb, M.S., et al. (2021). Profiling B cell immunodominance after SARS-CoV-2 infection reveals antibody evolution to non-neutralizing viral targets. Immunity 54, 1290–1303 e1297. 10.1016/j.immuni.2021.05.001.

39. Rosa, D., Saletti, G., De Gregorio, E., Zorat, F., Comar, C., D’Oro, U., Nuti, S., Houghton, M., Barnaba, V., Pozzato, G., and Abrignani, S. (2005). Activation of naive B lymphocytes via CD81, a pathogenetic mechanism for hepatitis C virus-associated B lymphocyte disorders. Proc. Natl. Acad. Sci. U. S. A. 102, 18544–18549. 10.1073/pnas.0509402102.

40. Pileri, P., Uematsu, Y., Campagnoli, S., Galli, G., Falugi, F., Petracca, R., Weiner, A.J., Houghton, M., Rosa, D., Grandi, G., and Abrignani, S. (1998). Binding of hepatitis C virus to CD81. Science 282, 938–941. 10.1126/science.282.5390.938.

41. Reyes, R.A., Batugedara, G., Dutta, P., Reers, A.B., Garza, R., Ssewanyana, I., Jagannathan, P., Feeney, M.E., Greenhouse, B., Bol, S., et al. (2023). Atypical B cells consist of subsets with distinct functional profiles. iScience 26, 108496. 10.1016/j.isci.2023.108496.

42. Holla, P., Dizon, B., Ambegaonkar, A.A., Rogel, N., Goldschmidt, E., Boddapati, A.K., Sohn, H., Sturdevant, D., Austin, J.W., Kardava, L., et al. (2021). Shared transcriptional profiles of atypical B cells suggest common drivers of expansion and function in malaria, HIV, and autoimmunity. Sci Adv 7. 10.1126/sciadv.abg8384.

43. Heim, M.H., and Thimme, R. (2014). Innate and adaptive immune responses in HCV infections. J. Hepatol. 61, S14–25. 10.1016/j.jhep.2014.06.035.

44. Snell, L.M., McGaha, T.L., and Brooks, D.G. (2017). Type I Interferon in Chronic Virus Infection and Cancer. Trends Immunol. 38, 542–557. 10.1016/j.it.2017.05.005.

45. Ticha, O., Moos, L., and Bekeredjian-Ding, I. (2021). Effects of long-term cryopreservation of PBMC on recovery of B cell subpopulations. J. Immunol. Methods 495, 113081. 10.1016/j.jim.2021.113081.

46. Nehar-Belaid, D., Hong, S., Marches, R., Chen, G., Bolisetty, M., Baisch, J., Walters, L., Punaro, M., Rossi, R.J., Chung, C.H., et al. (2020). Mapping systemic lupus erythematosus heterogeneity at the single-cell level. Nat. Immunol. 21, 1094–1106. 10.1038/s41590-020-0743-0.

47. Cooper, L., Xu, H., Polmear, J., Kealy, L., Szeto, C., Pang, E.S., Gupta, M., Kirn, A., Taylor, J.J., Jackson, K.J.L., et al. (2024). Type I interferons induce an epigenetically distinct memory B cell subset in chronic viral infection. Immunity 57, 1037–1055 e1036. 10.1016/j.immuni.2024.03.016.

48. Iwata, Y., Matsushita, T., Horikawa, M., Dilillo, D.J., Yanaba, K., Venturi, G.M., Szabolcs, P.M., Bernstein, S.H., Magro, C.M., Williams, A.D., et al. (2011). Characterization of a rare IL-10-competent B-cell subset in humans that parallels mouse regulatory B10 cells. Blood 117, 530–541. 10.1182/blood-2010-07-294249.

49. Rosser, E.C., and Mauri, C. (2015). Regulatory B cells: Origin, phenotype, and function. Immunity 42, 607–612. 10.1016/j.immuni.2015.04.005.

50. Yang, S.Y., Long, J., Huang, M.X., Luo, P.Y., Bian, Z.H., Xu, Y.F., Wang, C.B., Yang, S.H., Li, L., Selmi, C., et al. (2021). Characterization of organ-specific regulatory B cells using single-cell RNA sequencing. Front. Immunol. 12, 711980. 10.3389/fimmu.2021.711980.

51. Michee-Cospolite, M., Boudigou, M., Grasseau, A., Simon, Q., Mignen, O., Pers, J.O., Cornec, D., Le Pottier, L., and Hillion, S. (2022). Molecular mechanisms driving IL-10-producing B cells functions: STAT3 and c-MAF as underestimated central key regulators? Front. Immunol. 13, 818814. 10.3389/fimmu.2022.818814.

52. Ahsan, S., and Draghici, S. (2017). Identifying significantly impacted pathways and putative mechanisms with iPathwayGuide. Curr Protoc Bioinformatics 57, 7 1511-17 15 30. 10.1002/cpbi.24.

53. Brenner, D., Blaser, H., and Mak, T.W. (2015). Regulation of tumour necrosis factor signalling: live or let die. Nat. Rev. Immunol. 15, 362–374. 10.1038/nri3834.

54. Ticha, O., Slanina, P., Moos, L., Stichova, J., Vlkova, M., and Bekeredjian-Ding, I. (2021). TNFR2 expression is a hallmark of human memory B cells with suppressive function. Eur. J. Immunol. 51, 1195–1205. 10.1002/eji.202048988.

55. Ito, S., Kuromiya, K., Sekai, M., Sako, H., Sai, K., Morikawa, R., Mukai, Y., Ida, Y., Anzai, M., Ishikawa, S., et al. (2023). Accumulation of annexin A2 and S100A10 prevents apoptosis of apically delaminated, transformed epithelial cells. Proc. Natl. Acad. Sci. U. S. A. 120, e2307118120. 10.1073/pnas.2307118120.

56. Morris, L., Chen, X., Alam, M., Tomaras, G., Zhang, R., Marshall, D.J., Chen, B., Parks, R., Foulger, A., Jaeger, F., et al. (2011). Isolation of a human anti-HIV gp41 membrane proximal region neutralizing antibody by antigen-specific single B cell sorting. PLoS One 6, e23532. 10.1371/journal.pone.0023532.

57. Pushparaj, P., Nicoletto, A., Sheward, D.J., Das, H., Castro Dopico, X., Perez Vidakovics, L., Hanke, L., Chernyshev, M., Narang, S., Kim, S., et al. (2023). Immunoglobulin germline gene polymorphisms influence the function of SARS-CoV-2 neutralizing antibodies. Immunity 56, 193–206 e197. 10.1016/j.immuni.2022.12.005.

58. Sangesland, M., Torrents de la Pena, A., Boyoglu-Barnum, S., Ronsard, L., Mohamed, F.A.N., Moreno, T.B., Barnes, R.M., Rohrer, D., Lonberg, N., Ghebremichael, M., et al. (2022). Allelic polymorphism controls autoreactivity and vaccine elicitation of human broadly neutralizing antibodies against influenza virus. Immunity 55, 1693–1709 e1698. 10.1016/j.immuni.2022.07.006.

59. Shaffer, A.L., Shapiro-Shelef, M., Iwakoshi, N.N., Lee, A.H., Qian, S.B., Zhao, H., Yu, X., Yang, L., Tan, B.K., Rosenwald, A., et al. (2004). XBP1, downstream of Blimp-1, expands the secretory apparatus and other organelles, and increases protein synthesis in plasma cell differentiation. Immunity 21, 81–93. 10.1016/j.immuni.2004.06.010.

60. Johansen, F.E., Braathen, R., and Brandtzaeg, P. (2000). Role of J chain in secretory immunoglobulin formation. Scand. J. Immunol. 52, 240–248. 10.1046/j.1365-3083.2000.00790.x.

61. Salas, J.H., Urbanowicz, R.A., Guest, J.D., Frumento, N., Figueroa, A., Clark, K.E., Keck, Z., Cowton, V.M., Cole, S.J., Patel, A.H., et al. (2022). An Antigenically Diverse, Representative Panel of Envelope Glycoproteins for Hepatitis C Virus Vaccine Development. Gastroenterology 162, 562–574. 10.1053/j.gastro.2021.10.005.

62. Johnson, K., Green, P.K., and Ioannou, G.N. (2017). Implications of HCV RNA level at week 4 of direct antiviral treatments for hepatitis C. J. Viral Hepat. 24, 966–975. 10.1111/jvh.12731.

63. Mazouz, S., Boisvert, M., Shoukry, N.H., and Lamarre, D. (2018). Reversing immune dysfunction and liver damage after direct-acting antiviral treatment for hepatitis C. Can Liver J 1, 78–105. 10.3138/canlivj.1.2.007.

64. D’Ambrosio, R., Aghemo, A., Rumi, M.G., Ronchi, G., Donato, M.F., Paradis, V., Colombo, M., and Bedossa, P. (2012). A morphometric and immunohistochemical study to assess the benefit of a sustained virological response in hepatitis C virus patients with cirrhosis. Hepatology 56, 532–543. 10.1002/hep.25606.

65. Whitcomb, E., Choi, W.T., Jerome, K.R., Cook, L., Landis, C., Ahn, J., Te, H.S., Esfeh, J., Hanouneh, I.A., Rayhill, S.C., et al. (2017). Biopsy specimens from allograft liver contain histologic features of hepatitis C virus infection after virus eradication. Clin. Gastroenterol. Hepatol. 15, 1279–1285. 10.1016/j.cgh.2017.04.041.

66. van Loo, G., and Bertrand, M.J.M. (2023). Death by TNF: A road to inflammation. Nat. Rev. Immunol. 23, 289–303. 10.1038/s41577-022-00792-3.

67. Holbrook, J., Lara-Reyna, S., Jarosz-Griffiths, H., and McDermott, M. (2019). Tumour necrosis factor signalling in health and disease. F1000Res 8. 10.12688/f1000research.17023.1.

68. Heymann, F., and Tacke, F. (2016). Immunology in the liver-from homeostasis to disease. Nat. Rev. Gastroenterol. Hepatol. 13, 88–110. 10.1038/nrgastro.2015.200.

69. Portugal, S., Obeng-Adjei, N., Moir, S., Crompton, P.D., and Pierce, S.K. (2017). Atypical memory B cells in human chronic infectious diseases: An interim report. Cell. Immunol. 321, 18–25. 10.1016/j.cellimm.2017.07.003.

70. Nellore, A., Zumaquero, E., Scharer, C.D., Fucile, C.F., Tipton, C.M., King, R.G., Mi, T., Mousseau, B., Bradley, J.E., Zhou, F., et al. (2023). A transcriptionally distinct subset of influenza-specific effector memory B cells predicts long-lived antibody responses to vaccination in humans. Immunity 56, 847–863 e848. 10.1016/j.immuni.2023.03.001.

71. Chung, M.K.Y., Gong, L., Kwong, D.L., Lee, V.H., Lee, A.W., Guan, X.Y., Kam, N.W., and Dai, W. (2023). Functions of double-negative B cells in autoimmune diseases, infections, and cancers. EMBO Mol. Med. 15, e17341. 10.15252/emmm.202217341.

72. Sutton, H.J., Aye, R., Idris, A.H., Vistein, R., Nduati, E., Kai, O., Mwacharo, J., Li, X., Gao, X., Andrews, T.D., et al. (2021). Atypical B cells are part of an alternative lineage of B cells that participates in responses to vaccination and infection in humans. Cell Rep. 34, 108684. 10.1016/j.celrep.2020.108684.

73. Burchill, M.A., Salomon, M.P., Golden-Mason, L., Wieland, A., Maretti-Mira, A.C., Gale, M., Jr., and Rosen, H.R. (2021). Single-cell transcriptomic analyses of T cells in chronic HCV-infected patients dominated by DAA-induced interferon signaling changes. PLoS Pathog. 17, e1009799. 10.1371/journal.ppat.1009799.

74. Henning, A.N., Budeebazar, M., Boldbaatar, D., Yagaanbuyant, D., Duger, D., Batsukh, K., Zhou, H., Baumann, R., Allison, R.D., Alter, H.J., et al. (2023). Peripheral B cells from patients with hepatitis C virus-associated lymphoma exhibit clonal expansion and an anergic-like transcriptional profile. iScience 26, 105801. 10.1016/j.isci.2022.105801.

75. Lukhele, S., Boukhaled, G.M., and Brooks, D.G. (2019). Type I interferon signaling, regulation and gene stimulation in chronic virus infection. Semin. Immunol. 43, 101277. 10.1016/j.smim.2019.05.001.

76. Giordani, L., Sanchez, M., Libri, I., Quaranta, M.G., Mattioli, B., and Viora, M. (2009). IFN-α amplifies human naive B cell TLR-9-mediated activation and Ig production. J. Leukoc. Biol. 86, 261–271. 10.1189/jlb.0908560.

77. Ogega, C.O., Skinner, N.E., Schoenle, M.V., Wilcox, X.E., Frumento, N., Wright, D.A., Paul, H.T., Sinnis-Bourozikas, A., Clark, K.E., Figueroa, A., et al. (2024). Convergent evolution and targeting of diverse E2 epitopes by human broadly neutralizing antibodies are associated with HCV clearance. Immunity 57, 890–903 e896. 10.1016/j.immuni.2024.03.001.

78. Keck, Z.Y., Pierce, B.G., Lau, P., Lu, J., Wang, Y., Underwood, A., Bull, R.A., Prentoe, J., Velazquez-Moctezuma, R., Walker, M.R., et al. (2019). Broadly neutralizing antibodies from an individual that naturally cleared multiple hepatitis C virus infections uncover molecular determinants for E2 targeting and vaccine design. PLoS Pathog. 15, e1007772. 10.1371/journal.ppat.1007772.

79. Chen, F., Nagy, K., Chavez, D., Willis, S., McBride, R., Giang, E., Honda, A., Bukh, J., Ordoukhanian, P., Zhu, J., et al. (2020). Antibody responses to immunization with HCV envelope glycoproteins as a baseline for B-cell-based vaccine development. Gastroenterology 158, 1058–1071 e1056. 10.1053/j.gastro.2019.11.282.

80. Chen, F., Tzarum, N., Lin, X., Giang, E., Velazquez-Moctezuma, R., Augestad, E.H., Nagy, K., He, L., Hernandez, M., Fouch, M.E., et al. (2021). Functional convergence of a germline-encoded neutralizing antibody response in rhesus macaques immunized with HCV envelope glycoproteins. Immunity 54, 781–796 e784. 10.1016/j.immuni.2021.02.013.

81. Underwood, A.P., Gupta, M., Wu, B.R., Eltahla, A.A., Boo, I., Wang, J.J., Agapiou, D., Abayasingam, A., Reynaldi, A., Keoshkerian, E., et al. (2024). B-cell characteristics define HCV reinfection outcome. J. Hepatol. 81, 415–428. 10.1016/j.jhep.2024.04.004.

82. Fu, Y., Yip, A., Seah, P.G., Blasco, F., Shi, P.Y., and Herve, M. (2014). Modulation of inflammation and pathology during dengue virus infection by p38 MAPK inhibitor SB203580. Antiviral Res. 110, 151–157. 10.1016/j.antiviral.2014.08.004.

83. Sreekanth, G.P., Chuncharunee, A., Sirimontaporn, A., Panaampon, J., Noisakran, S., Yenchitsomanus, P.T., and Limjindaporn, T. (2016). SB203580 modulates p38 MAPK signaling and Dengue virus-induced liver injury by reducing MAPKAPK2, HSP27, and ATF2 phosphorylation. PLoS One 11, e0149486. 10.1371/journal.pone.0149486.

84. Madkour, M.M., Anbar, H.S., and El-Gamal, M.I. (2021). Current status and future prospects of p38alpha/MAPK14 kinase and its inhibitors. Eur. J. Med. Chem. 213, 113216. 10.1016/j.ejmech.2021.113216.

85. Barnes, E., Cooke, G.S., Lauer, G.M., and Chung, R.T. (2023). Implementation of a controlled human infection model for evaluation of HCV vaccine candidates. Hepatology 77, 1757–1772. 10.1002/hep.32632.

86. Feld, J.J., Bruneau, J., Dore, G.J., Ghany, M.G., Hansen, B., Sulkowski, M., and Thomas, D.L. (2023). Controlled human infection model for hepatitis C virus vaccine development: Trial design considerations. Clin. Infect. Dis. 77, S262–S269. 10.1093/cid/ciad362.

87. Liang, T.J., Law, J.L.M., Pietschmann, T., Ray, S.C., Bukh, J., Bull, R., Chung, R.T., Tyrrell, D.L., Houghton, M., and Rice, C.M. (2023). Challenge inoculum for hepatitis C virus controlled human infection model. Clin. Infect. Dis. 77, S257–S261. 10.1093/cid/ciad336.

88. Borrego-Yaniz, G., Terron-Camero, L.C., Kerick, M., Andres-Leon, E., and Martin, J. (2024). A holistic approach to understanding immune-mediated inflammatory diseases: bioinformatic tools to integrate omics data. Comput Struct Biotechnol J 23, 96–105. 10.1016/j.csbj.2023.11.045.

89. Wang, R., Suzuki, S., Guest, J.D., Heller, B., Almeda, M., Andrianov, A.K., Marin, A., Mariuzza, R.A., Keck, Z.Y., Foung, S.K.H., et al. (2022). Induction of broadly neutralizing antibodies using a secreted form of the hepatitis C virus E1E2 heterodimer as a vaccine candidate. Proc. Natl. Acad. Sci. U. S. A. 119, e2112008119. 10.1073/pnas.2112008119.

90. Zhou, P., Song, G., Liu, H., Yuan, M., He, W.T., Beutler, N., Zhu, X., Tse, L.V., Martinez, D.R., Schafer, A., et al. (2023). Broadly neutralizing anti-S2 antibodies protect against all three human betacoronaviruses that cause deadly disease. Immunity 56, 669–686 e667. 10.1016/j.immuni.2023.02.005.

91. Weaver, G.C., Villar, R.F., Kanekiyo, M., Nabel, G.J., Mascola, J.R., and Lingwood, D. (2016). In vitro reconstitution of B cell receptor-antigen interactions to evaluate potential vaccine candidates. Nat. Protoc. 11, 193–213. 10.1038/nprot.2016.009.

92. Zheng, G.X., Terry, J.M., Belgrader, P., Ryvkin, P., Bent, Z.W., Wilson, R., Ziraldo, S.B., Wheeler, T.D., McDermott, G.P., Zhu, J., et al. (2017). Massively parallel digital transcriptional profiling of single cells. Nat Commun 8, 14049. 10.1038/ncomms14049.

93. Gupta, N.T., Vander Heiden, J.A., Uduman, M., Gadala-Maria, D., Yaari, G., and Kleinstein, S.H. (2015). Change-O: a toolkit for analyzing large-scale B cell immunoglobulin repertoire sequencing data. Bioinformatics 31, 3356–3358. 10.1093/bioinformatics/btv359.

94. McKenna, A., Hanna, M., Banks, E., Sivachenko, A., Cibulskis, K., Kernytsky, A., Garimella, K., Altshuler, D., Gabriel, S., Daly, M., and DePristo, M.A. (2010). The genome analysis toolkit: A MapReduce framework for analyzing next-generation DNA sequencing data. Genome Res. 20, 1297–1303. 10.1101/gr.107524.110.

95. Huang, X., and Huang, Y. (2021). Cellsnp-lite: An efficient tool for genotyping single cells. Bioinformatics 37, 4569–4571. 10.1093/bioinformatics/btab358.

96. Major, M., and Law, M. (2019). Detection of antibodies to HCV E1E2 by lectin-capture ELISA. Methods Mol. Biol. 1911, 421–432. 10.1007/978-1-4939-8976-8_28.

97. Kalemera, M.D., Capella-Pujol, J., Chumbe, A., Underwood, A., Bull, R.A., Schinkel, J., Sliepen, K., and Grove, J. (2021). Optimized cell systems for the investigation of hepatitis C virus E1E2 glycoproteins. J. Gen. Virol. 102. 10.1099/jgv.0.001512.

98. Bailey, J.R., Urbanowicz, R.A., Ball, J.K., Law, M., and Foung, S.K.H. (2019). Standardized method for the study of antibody neutralization of HCV pseudoparticles (HCVpp). Methods Mol. Biol. 1911, 441–450. 10.1007/978-1-4939-8976-8_30.

